# Dynamics of the N-terminal domain of SARS-CoV-2 nucleocapsid protein drives dsRNA melting in a counterintuitive tweezer-like mechanism

**DOI:** 10.1101/2020.08.24.264465

**Authors:** Ícaro P. Caruso, Karoline Sanches, Andrea T. Da Poian, Anderson S. Pinheiro, Fabio C. L. Almeida

## Abstract

The N protein of betacoronaviruses is responsible for nucleocapsid assembly and other essential regulatory functions. Its N-terminal domain (NTD) interacts and melts the double-stranded transcriptional regulatory sequences (dsTRS), regulating the discontinuous subgenome transcription process. Here, we used molecular dynamics (MD) simulations to study the binding of SARS-CoV-2 N-NTD to non-specific (NS) and TRS dsRNAs. We probed dsRNAs’ Watson and Crick (WC) base-pairing over 25 replicas of 100 ns MD simulations, showing that only one N-NTD of dimeric N is enough to destabilize dsRNAs, initiating melting. N-NTD dsRNA destabilizing activity was more efficient for dsTRS than dsNS. N-NTD dynamics, especially a tweezer-like motion of β2-β3 and 2-β5 loops, played a key role in WC base-pairing destabilization. Based on experimental information available in the literature, we constructed kinetics models for N-NTD-mediated dsRNA melting. Our results support a 1:1 stoichiometry (N-NTD:dsRNA), matching MD simulations and raising different possibilities for N-NTD action: (i) two N-NTDs of dimeric N would act independently, increasing efficiency; (ii) two N-NTDs of dimeric N would bind to two different RNA sites, bridging distant regions of the genome; and (iii) monomeric N would be active, opening up the possibility of a regulatory dissociation event.

**IMPORTANCE:** Coronaviruses are among the largest positive-sense RNA viruses. They display a unique discontinous transcription mechanism, involving N protein as a major player. The N-NTD promote the dsRNA melting releasing the nascent sense negative strand via a poorly known mechanism of action. It specifically recognizes the body TRS conserved RNA motif located at the 5’ end of each ORF. N protein has the ability to transfer the nascent RNA strand to the leader TRS. The mechanism is essential and one single mutation at the RNA binding site of the N-NTD impairs the viral replication. Here, we describe a counterintuitive mechanism of action of N-NTD based on molecular dynamics simulation and kinetic modelling of the experimental melting activity of N-NTD. This data impacts directly in the understanding of the way N protein acts in the cell and will guide future experiments.

## INTRODUCTION

The recent pandemic of Severe Acute Respiratory Syndrome Coronavirus 2 (SARS-CoV-2), the causative agent of Coronavirus Disease 2019 (COVID-19), has become a global health emergency (1, 2). SARS-CoV-2 is an enveloped virus containing a large non-segmented positive-sense single-stranded RNA genome, belonging to the *Coronaviridae* family (3, 4). The 5’ two-thirds of the coronaviruses’ genome, corresponding to ORF1a/b, is translated into two polyproteins (pp1a and pp1ab) that are proteolytically processed into sixteen nonstructural proteins (NSPs) (5). These NSPs assemble in the viral replicase-transcriptase complex (RTC) at the endoplasmic reticulum membrane, being responsible for genome replication and transcription (6). Conversely, the 3’ one-third of the genome is translated into accessory proteins as well as the four structural proteins, spike (S), membrane (M), envelope (E), and nucleocapsid (N), through a unique process of subgenomic mRNA (sgmRNA) transcription (7, 8).

N is one of the most abundant viral proteins in the infected cell. It is a 46-kDa multifunctional RNA-binding protein that drives the viral RNA packaging into a helical nucleocapsid (9). In addition, N localizes at the RTC at early stages of infection and plays a central role in the regulation of RNA synthesis (10–12). It is composed of two functionally distinct folded domains, which are interspersed by an intrinsically disordered linker region enriched in arginine and serine residues. Both the two domains and the linker region contribute individually to RNA binding (13). The N-terminal domain (NTD) has been shown to interact with regulatory RNA sequences during subgenome transcription, whereas the C-terminal domain (CTD) is responsible for N protein dimerization, which is crucial for nucleocapsid assembly (14, 15). The recently reported solution structure of SARS-CoV-2 N-NTD reveals a right hand-like fold, composed of a five-stranded central β-sheet flanked by two short α-helices, arranged in a β4-β2-β3-β1-β5 topology (16). The β-sheet core is referred to as the hand’s palm, while the long β2-β3 hairpin, mostly composed of basic amino acid residues, corresponds to the basic finger. The positively-charged cleft between the basic finger and the palm has been suggested as a putative RNA binding site (16).

Genome replication is a continuous process in coronaviruses. In contrast, transcription is discontinuous and involves the production of sgmRNAs (17). Regulation of sgmRNA synthesis is dependent on transcriptional regulatory sequences (TRSs) located either at the 5’ end of the positive-strand RNA genome, known as the leader TRS (TRS-L), or at the 5’ end of each viral gene coding for structural and accessory proteins, called the body TRS (TRS-B). TRS-L and TRS-B share an identical core sequence, which allows for a template switch during sgmRNA synthesis. Once the TRS-B has been copied, the nascent negative-strand RNA is transferred to TRS-L and transcription is terminated (17, 18). Multiple well-orchestrated factors, including TRS secondary structure, RNA-RNA and RNA-protein interactions, influence sgmRNA transcription (17). Coronaviruses’ N-NTD specifically interacts with the TRS and efficiently melts a TRS-cTRS RNA duplex, facilitating template switch and playing a pivotal role in the regulation of discontinuous transcription (10, 16, 17, 19). Despite its relevance for the viral replication cycle, the molecular basis underlying the specificity of interaction of SARS-CoV-2 N-NTD with the TRS sequence remains elusive. Thus, understanding the mechanism by which SARS-CoV-2 N-NTD specifically recognizes TRS RNA at atomic detail is paramount for the rational development of new antiviral strategies.

Here, we present a hypothesis for the molecular mechanism of dsRNA melting activity of SARS-CoV-2 N-NTD. We showed by molecular dynamics (MD) simulations (25 replicas of 100 ns) that N-NTD destabilizes dsRNA’s Watson and Crick base-pairing by dropping down intramolecular hydrogen bonds and perturbing the local rigid-body geometric parameters of dsRNA. The destabilization is more significant for TRS than for a non-specific (NS) dsRNA sequence. Moreover, a tweezer-like motion between β2-β3 and α2-β5 loops of N-NTD seems to be a key dynamic feature for selectivity and, consequently, dsRNA melting activity. We also constructed kinetic models for characterizing the melting activity of the dimeric N protein assuming 1:1 and 2:1 (N-NTD:dsRNA) stoichiometries, revealing that only one N-NTD is enough for dsRNA melting.

## RESULTS

### Structural models of the N-NTD:dsRNA complexes and their validation from molecular dynamics simulations

We calculated the structural model of the N-NTD:dsTRS complex based on the experimental data for the N-NTD interaction with a non-specific dsRNA (5’–CACUGAC– 3’) (dsNS) (16) using the HADDOCK 2.2 server (20). The structural restraints of the N-NTD:dsNS complex were defined from CSPs titration performed by Dinesh et al. (2020) (16). The lowest-energy structure of the N-NTD:dsNS complex from the cluster with the lowest HADDOCK score (fraction common contacts = 0.8 ± 0.1 Å, interface-RMSD = 1.0 ± 0.6 Å, and ligand-RMSD = 2.2 ± 1.2 Å) was used to mutate the dsNS molecule to obtain the TRS sequence (5’–UCUAAAC–3’) and, therefore, to generate the N-NTD:dsTRS complex structure. Figure 1A shows the structural model of the N-NTD:dsTRS complex, in which the TRS RNA is inserted in a cleft located between the large protruding β2-β3 loop, named finger, and the central β-sheet of N-NTD, referred to as the palm. Analysis of the electrostatic surface potential of N-NTD revealed that the dsRNA-binding pocket is positively charged, with the finger being the highest charged region (Figure S1). This result is consistent with the charge complementarity of the nucleic acid phosphate groups that exhibit negative charge. It is worth mentioning that the orientation of the TRS sense strand in the complex model is in agreement with experimental results described by Keane et al. (2012) for the Murine Hepatitis Virus structural homolog of N-NTD (21), in which the 5’-end of the sense strand binds close to β4-β2-β3 and the 3’-end binds next to β1-β5 (Figure 1A).

**Figure 1.**
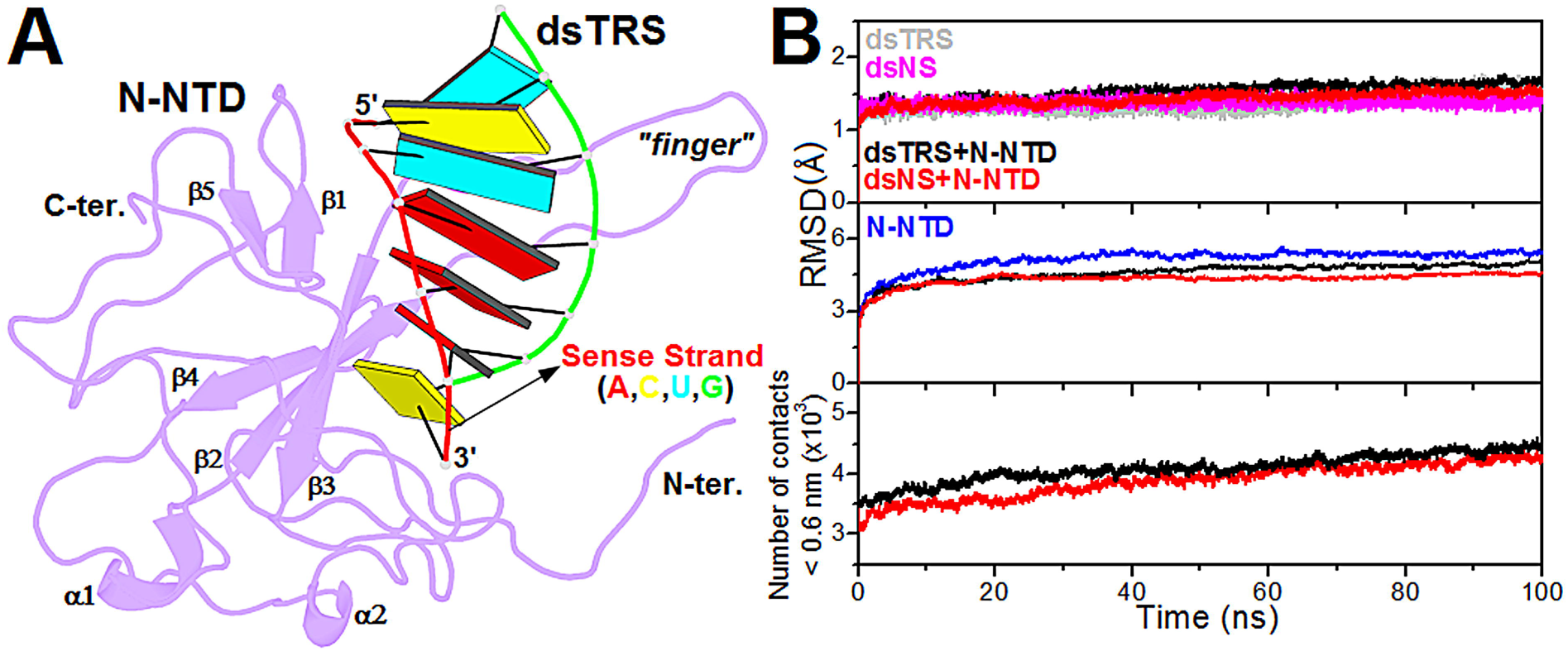
Structural model of the N-NTD:dsRNA complex and its validation from molecular dynamics simulations. (A) Structural model of the N-NTD:dsTRS complex determined by molecular docking calculations and mutation of dsNS nucleotide sequence. N-NTD is presented as purple cartoon and dsTRS is denoted as a ribbon model with base pairing as colored rectangles. The color of the rectangles corresponds to the nitrogenous base of the dsRNA sense strand, namely A: red, C: yellow, U: cyan, and G: green. The large protruding β2-β3 loop is referred to as the finger. (B) Average RMSD values for dsNS and dsTRS in their free and N-NTD-bound states (*top*), average RMSD values for N-NTD in its free and dsRNA-bound state (*middle*), and average number of contacts between N-NTD and dsRNA atoms (distance < 0.6 nm) (*bottom*). The average values correspond to 25 MD simulations with the same starting point.

We performed 25 calculations of 100 ns molecular dynamics (MD) simulations to investigate the stability of the structural models of N-NTD in complex with either dsNS or dsTRS, as well as each of the biomolecules separately (dsNS, dsTRS, and N-NTD). Figure 1B (*top*) shows the average RMSD values for the backbone atoms (C5’, C4’, C3’, O3’, P, and O5’) of dsNS and dsTRS in their free and N-NTD-bound states, which were significantly stable along the 100 ns MD simulations. Similar results are observed for the average RMSD values for free and dsRNA-bound N-NTD over the simulations (Figure 1B – *middle*). Evaluation of the average number of contacts between N-NTD and dsRNAs (distance < 0.6 nm) revealed that dsNS and dsTRS are in close interaction with N-NTD throughout the 100 ns MD simulations (Figure 1B – *bottom*). These parameters (RMSD and contacts) validate the robustness and stability of the structural models generated for the N-NTD:dsRNA complexes as well as the molecular structures of the investigated biomolecules (dsNS, dsTRS, and N-NTD). The non-averaged values of the analyzed parameters for each of the 25 MD simulations are provided in Supplementary Material (Figure S2–S10).

### Stability of dsNS and dsTRS base-pairing upon N-NTD binding

To estimate the stability of the Watson-Crick (WC) base-pairing of dsNS and dsTRS complexed with N-NTD, we evaluated the intramolecular hydrogen bonds formed between sense and anti-sense strands of the dsRNA bound to N-NTD. In addition, the hydrogen bonds of the free RNA molecules were investigated as a control parameter. Figure 2 shows the number of intramolecular hydrogen bonds for 25 replicas of MD simulations of free and N-NTD-bound dsTRS. It is possible to note that the score profile of intramolecular hydrogen bonds for free dsTRS (Figure 2A) was different than that of N-NTD-bound dsTRS (Figure 2B). This difference is mainly due to a considerable reduction in the number of intramolecular hydrogen bonds in at least 4 replicas of the set of 25 MD simulations (runs 5, 8, 17, and 25), suggesting that dsTRS WC base-pairing was destabilized by interaction with N-NTD. On the other hand, the score profile of intramolecular hydrogen bonds for free dsNS (Figure 2C) was quite similar to its N-NTD-bound state (Figure 2D).

**Figure 2.**
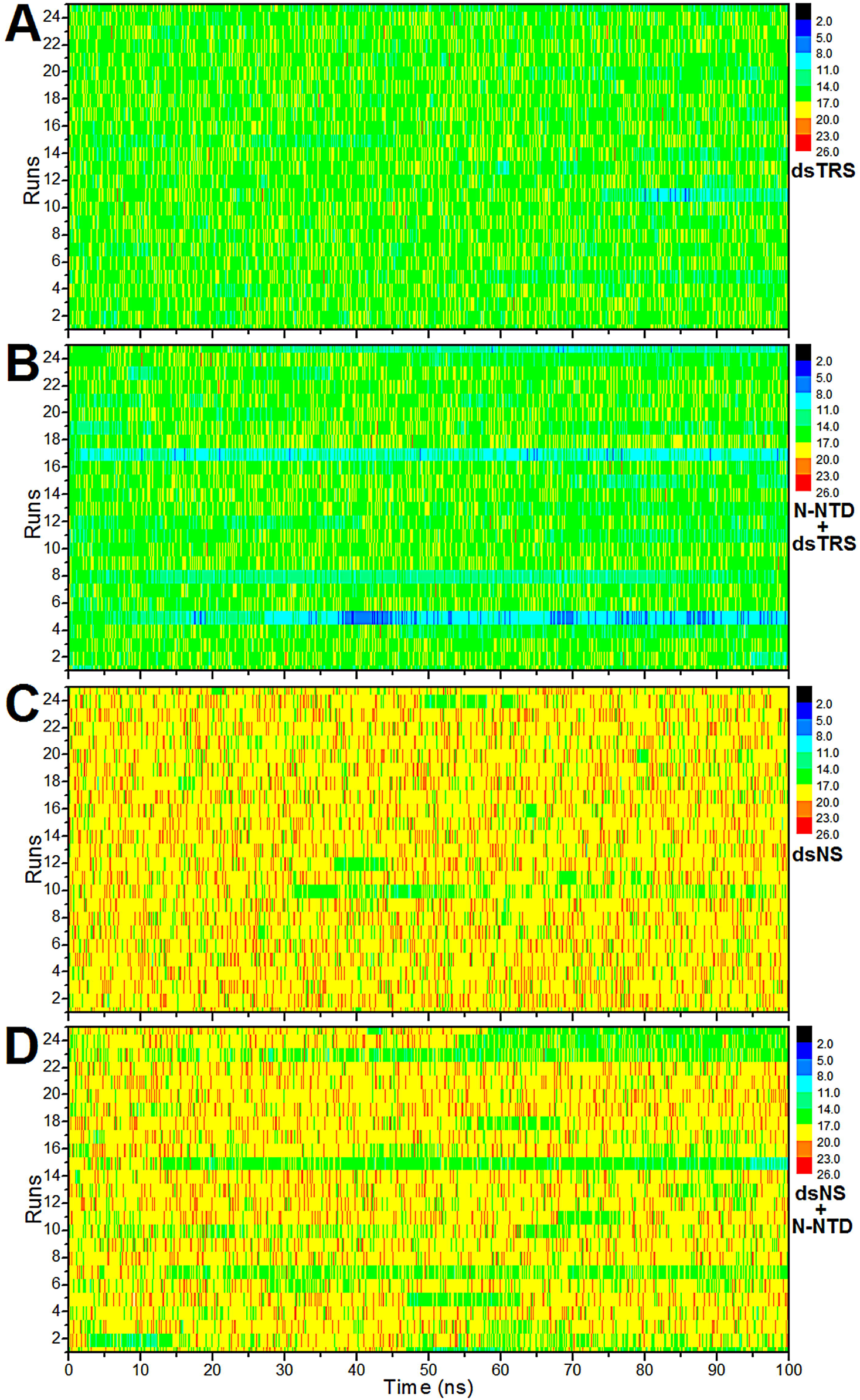
Stability of the Watson-Crick base-pairing via intramolecular hydrogen bonds of dsRNAs. The number of intramolecular hydrogen bonds formed between the sense and anti-sense strands of dsNS and dsTRS in their free states (A, C) and in complex with N-NTD (B, D) over the 100 ns simulations for the 25 MD replicas (runs). The color bar denotes the correspondence between the color code and the number of intramolecular hydrogen bonds.

We also performed a quantitative analysis of WC base-pairing by calculating the average number of intramolecular hydrogen bonds throughout the 100 ns MD simulations for each of the 25 replicas. For free dsRNAs, the average numbers of intramolecular hydrogen bonds were constant for the 25 runs (Figure 3A) with an overall average value of 18.2 ± 0.2 and 15.5 ± 0.3 for dsNS and dsTRS, respectively. It is worth noting that the expected value of WC hydrogen bonds for dsNS and dsTRS are 18 and 16, respectively. This confirms the consistency of the force field used to describe the studied molecular system. For the N-NTD-bound dsRNAs, we observed a decrease in the overall average number of intramolecular hydrogen bonds for both RNA ligands, accompanied by an increase in respective standard deviations. This increase in standard deviation is due to a significant reduction in the average number of intramolecular hydrogen bonds of particular replicas, specifically 4 for dsTRS (runs 5, 8, 17, and 25) and 2 for dsNS (runs 15 and 23) (Figure 3A). The N-NTD-induced reduction in the number of intramolecular hydrogen bonds between the sense and anti-sense strands of dsRNAs was more pronounced for dsTRS than dsNS, as shown by the analysis of score profile in Figure 2.

**Figure 3.**
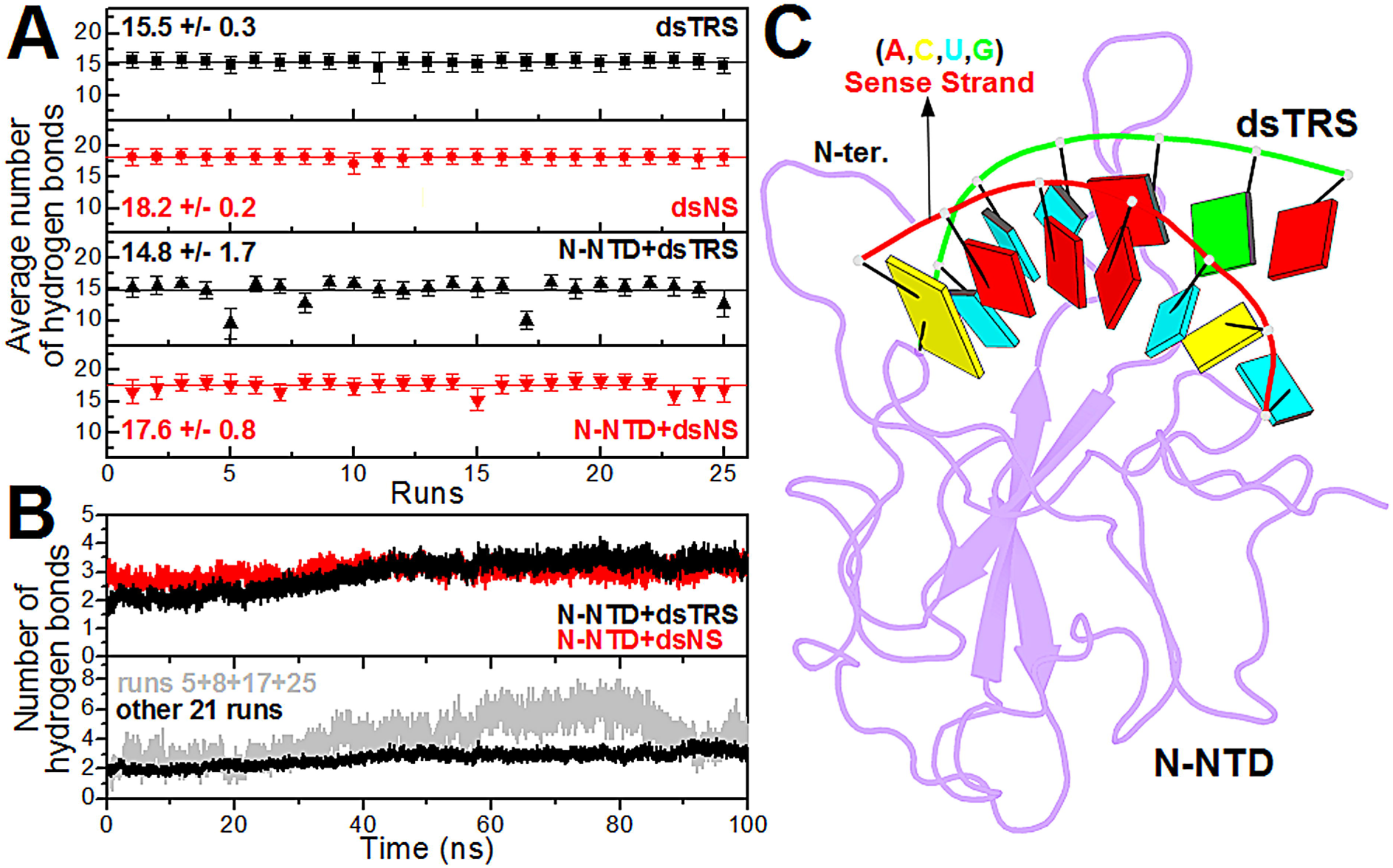
Analysis of the intramolecular (dsRNAs) and intermolecular (N-NTD:dsRNAs) hydrogen bonds. (A) The average number of intramolecular hydrogen bonds between the sense and anti-sense strands of dsNS (red) and dsTRS (black) in their free states (squares and circles) and in complex with N-NTD (up and down triangles, respectively) for each of the 25 replicas of 100 ns MD simulation. The black and red solid lines denote the overall average values for the 25 runs, which are also presented numerically with their respective standard deviations. (B) Average number of intermolecular hydrogen bonds between the nitrogenous bases of dsRNA (dsTRS in black and dsNS in red) and N-NTD for the 25 replicas along the 100 ns MD simulations (*top*). Average number of intermolecular hydrogen bonds for runs 5, 8, 17, and 25 (light gray) and the other 21 replicas (black) are shown as a function of simulation time (*bottom*). (C) Structural model of the N-NTD:dsTRS complex representative of the MD simulation for run 5. The protein is shown in purple cartoon and dsTRS is denoted as a ribbon model with nitrogenous bases and base-pairing as colored squares and rectangles, respectively. The color of the squares corresponds to the type of nitrogenous base, namely A: red, C: yellow, U: cyan, and G: green, while the rectangles refer to the nitrogenous base color of the dsRNA sense strand.

In contrast to the decrease in the number of intramolecular hydrogen bonds between the sense and anti-sense strands of dsRNAs (WC base-pairing) due to N-NTD binding, we observed an increase in the average number of intermolecular hydrogen bonds formed between the nitrogenous bases of dsTRS and N-NTD (protein-RNA interaction) along the 100 ns MD simulation, whereas for dsNS, this average value was constant (Figure 3B – *top*). It is noteworthy that dsNS has more hydrogen bond-forming sites (acceptor and donor) than dsTRS and, in spite of that, the average number of intermolecular hydrogen bonds for the N-NTD:dsTRS complex is higher after 50 ns simulation. Figure 3B (*bottom*) shows that the average number of intermolecular hydrogen bonds for runs 5, 8, 17, and 25 increased significantly with respect to the average value of the other replicas (21 runs in total). The non-averaged values of the intermolecular hydrogen bonds for each of the 25 MD simulations are provided in Supplementary Material (Figure S11). The drastic drop in the number of intramolecular hydrogen bonds for dsTRS in these 4 replicas is a consequence of the destabilization of the RNA duplex. Note that a complete break of WC base-pairing was observed in run 5 (Figure 3C), whereas a similar behavior with partial breaks occurred for dsTRS in runs 8, 17, and 25 (Figure S12). The structural model of the N-NTD:dsTRS complex presented in Figure 3C suggests a hypothesis for the mechanism of action of N-NTD in which only one domain is capable of destabilizing the RNA duplex, possibly leading to its dissociation and ultimately release of the RNA single strands.

To further understand the stability of the Watson-Crick base-pairing for N-NTD-bound dsNS and dsTRS, we used the *do_x3dna* tool (22) along with the 3DNA package (23) to analyze the local base-pair parameters (angles: buckle, opening, and propeller; distances: stretch, stagger, and shear) from the MD simulations. Figure 4 shows the population distributions of these local base-pair parameters for runs 5, 8, 17, and 25 (for dsTRS), and runs 15 and 23 (for dsNS), both free and complexed with N-NTD, as well as the difference between the distributions of the free and N-NTD-bound states. These replicas were selected based on the results presented in Figure 3A, since their average numbers of intramolecular hydrogen bonds were significantly lower than the overall average values. From Figure 4, it is clear that N-NTD perturbs the population distributions of the local base-pair parameters of dsRNAs, most notably for dsTRS.

**Figure 4.**
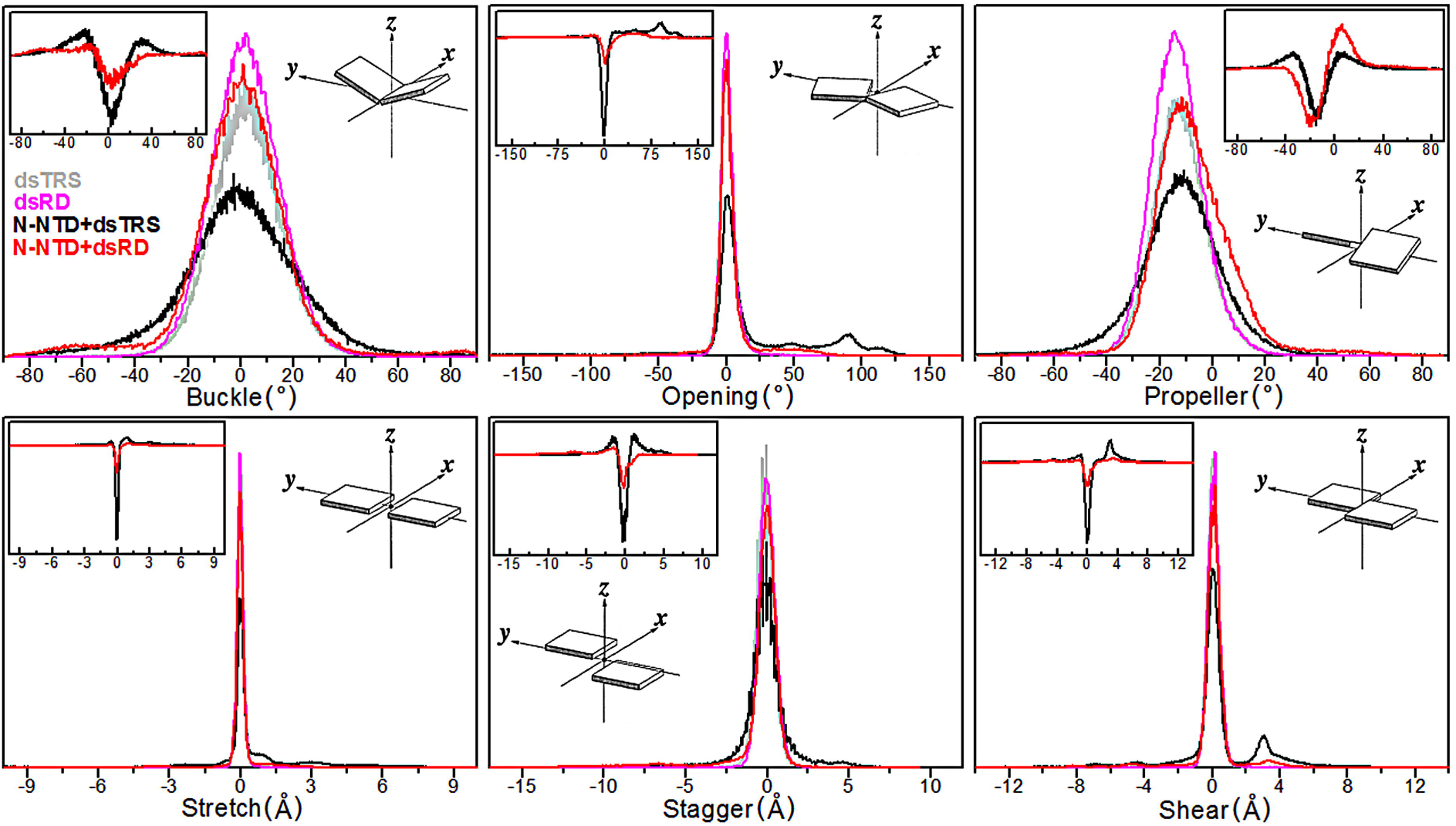
Population distributions of the local base-pair parameters. Population distributions of local base-pair parameters (angles: buckle, opening, and propeller; distances: stretch, stagger, and shear) for runs 5, 8, 17, and 25 of dsTRS and runs 15 and 23 of dsNS in their free form (dsTRS in light gray and dsNS in magenta, respectively) and complexed with N-NTD (N-NTD+dsTRS in black and N-NTD+dsNS in red). The plot insets correspond to the difference between the population distributions of N-NTD-bound dsRNA minus its free state for dsNS (red) and dsTRS (black). The scheme insets illustrate the geometrical definition of each local base-pair parameter (39).

The distribution of dsTRS buckle angles revealed a reduction in the population at ∼0° (base-pairing planarity) and increase in subpopulations at ∼±30° due to the interaction with N-NTD. This can be clearly seen by the difference in distributions between the N-NTD-bound and free states (inset in buckle plot of Figure 4). A similar but less intense effect was observed for the population distribution of dsNS buckle angles. For the opening angles, one can note a higher perturbation of population distribution for dsTRS than for dsNS upon binding to N-NTD. For dsTRS, the opening angle population at ∼0° (base-pairing closure) decreased significantly while subpopulations emerged for angles higher than 50°, remarkably at ∼90°. The distribution of the propeller angles showed a reduction in the equilibrium populations at –12.6° and –13.8° for dsTRS and dsNS, respectively, after interaction with N-NTD, with an increase in subpopulations around 0° (less twist), which was significantly larger for dsNS. However, we observed an extra subpopulation of propeller angles at approximately –30° for dsTRS after binding to N-NTD, which was not seen for dsNS (see the inset in propeller plot in Figure 4).

Investigation of the stretch, stagger, and shear distances for dsNS and dsTRS showed that the equilibrium population at ∼0 Å decreased for both dsRNAs as a result of N-NTD binding. However, this reduction is more drastic for dsTRS than dsNS, as can be seen in the inset for the respective plots in Figure 4. In addition to this reduction effect, we also verified that N-NTD-bound dsTRS exhibited clear subpopulations at ∼1, ∼±1.5, and ∼3 Å for the stretch, stagger, and shear distances, respectively.

N-NTD-induced perturbations in the population distributions of angle and distance base-pair parameters (buckle, opening, propeller, stretch, stagger, and shear) of dsRNAs for the selected replicates (runs 5, 8, 17, and 25 for dsTRS, and runs 15 and 23 for dsNS) indicate that both dsRNAs suffered WC base-pairing destabilization upon N-NTD binding. However, this destabilization effect is more evident for the N-NTD:dsTRS complex, since the above analysis of angle and distance parameters suggests an impairment of base-pairing planarity accompanied by an increase in the separation between the nitrogenous bases of the complementary dsRNA strands upon N-NTD binding. This result agrees well with the analysis of the intramolecular hydrogen bonds formed between the sense and anti-sense dsRNA strands (see Figure 3A). It is worth mentioning that, even though base-pairing destabilization was more pronounced for dsTRS than dsNS, dsNS suffered a greater reduction in the RNA duplex twist, as suggested by the N-NTD-induced perturbation of the propeller angles.

We also analyzed the population distributions of the local base-pair parameters for the 25 replicas of dsNS and dsTRS in their free and N-NTD-bound states. Figure S13 shows that the WC base-pairing perturbations observed in runs 5, 8, 17, and 25 for dsTRS due to N-NTD binding was also seen for the fully unbiased distribution generated from the 25 replicas. Nevertheless, an opposite effect occurred for dsNS, for which the angle and distance parameters exhibited characteristics of stability and/or slight fluctuations around the equilibrium population.

### Conformational flexibility of free and dsRNA-bound N-NTD

To further understand how dsRNA binding changes the conformational dynamics of N-NTD, we concatenated the last 50 ns (stable RMSD values) of the 25 replicas for both free and dsRNA-bound N-NTD and performed an analysis of root-mean-square fluctuation (RMSF) and principal component analysis (PCA) of the MD trajectories (Figure 5). Figure 5A shows that both free and dsRNA-bound N-NTD exhibited significantly increased values of RMSF for residues in the N- and C-terminal regions as well as the β2-β3 loop (finger), suggesting large conformational flexibility. In addition to these regions, the N-terminal portion of the β1-1 loop (residues 58–65) and β3-β4 loop are especially noteworthy. These loops displayed an increase in their dynamics in the N-NTD:dsTRS complex when compared to free N-NTD, even though they are not directly involved in the interaction. A gain in conformational flexibility can also be noted for the basic finger when N-NTD is complexed with dsTRS. In general, we observed an increase in flexibility of N-NTD loop regions when bound to dsTRS. Remarkably, conformational dynamics of the N-NTD:dsNS complex was similar to that of the free state, with the exception of the N-terminal region and basic finger, in which conformational dynamics decreased upon dsNS binding.

**Figure 5.**
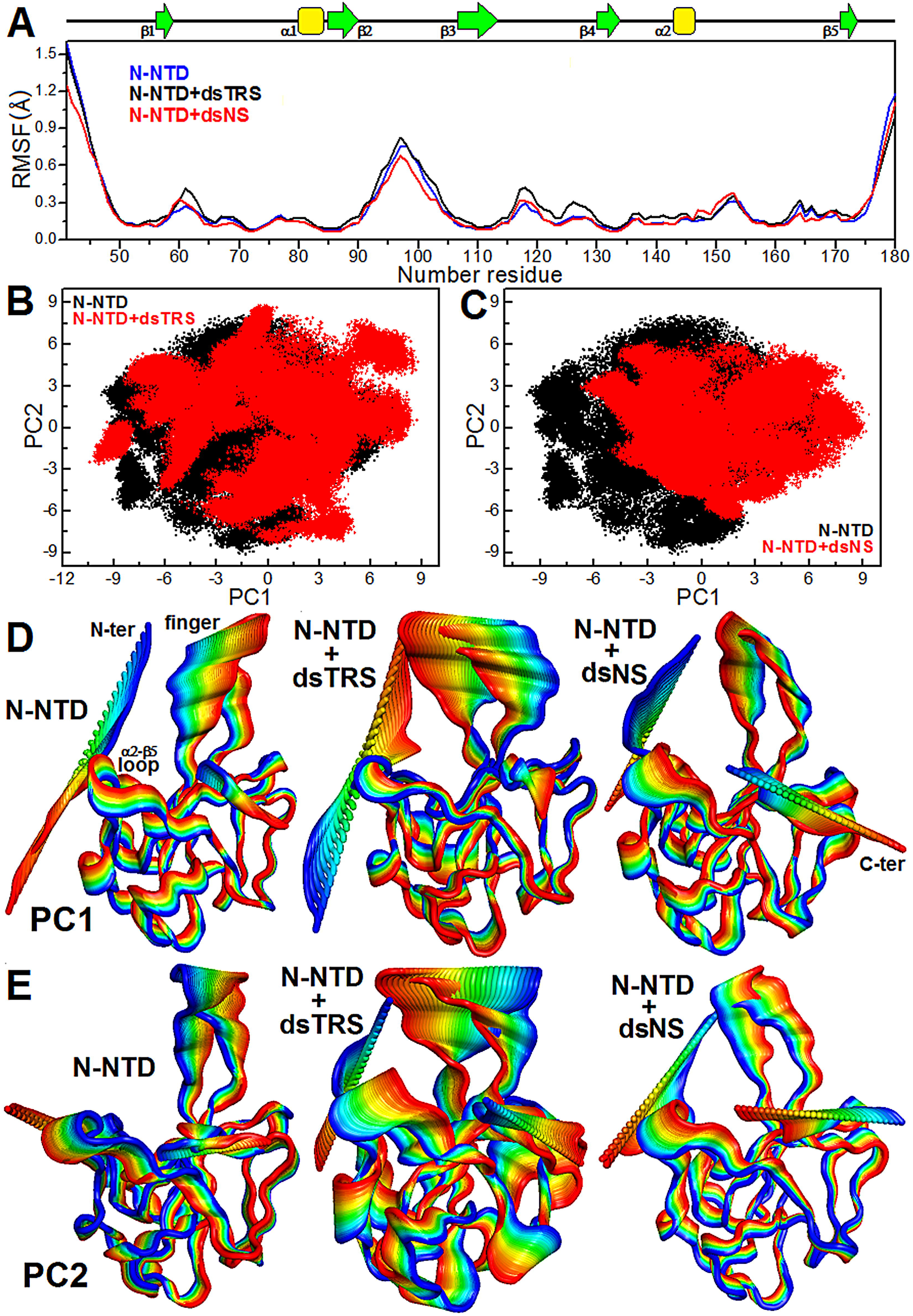
Analysis of N-NTD conformational flexibility in its free and dsRNA-bound states. (A) RMSF values as a function of residue number for N-NTD in its free state (blue line) and complexed with either dsTRS (black) or dsNS (red). The secondary structures along the sequence are indicated at the top. (B, C) PCA scatter plots PC1 and PC2 for free N-NTD (black dots) and for N-NTD complexed with either dsTRS (red, B) or dsNS (red, C) dsRNAs. (D, E) Motions filtered from the eigenvectors of PC1 (D) and PC2 (E) for the dynamics data of N-NTD in its free form and complexed with either dsTRS or dsNS. The motion direction is indicated by the color variation from blue to red. To perform the RMSF and PCA calculations, the last 50 ns of trajectories of the 25 replicates were concatenated for each of the molecular systems (free or dsRNA-bound N-NTD), resulting in MD simulations of 1.25 μs.

The PCA scatter plot generated for free and dsRNA-bound N-NTD revealed a significant difference between the free domain and the complexes, as evident from the characteristic structures plotted along the direction of the first and second principal components (PC1 and PC2, respectively). 25 replicas of the MD simulations using different seeds of the random number generator provided a great exploration of the conformational space for free and dsRNA-bound N-NTD, resulting in trajectories of 1.25 μs. Despite the already wide conformational space of N-NTD, dsTRS interaction made it even wider (Figure 5B), while dsNS binding made it more constrained (Figure 5C). An investigation of the motions filtered from the eigenvectors of PC1 and PC2 revealed that dsTRS-bound N-NTD exhibited the largest conformational dynamics when compared to free and dsNS-bound N-NTD, which were similar (Figure 5D and 5E). We highlight that the most evident motions took place in the N- and C-termini as well as the basic finger (β2-β3 loop) for both free and dsRNA-bound N-NTD. However, the eigenvectors of PC1 and PC2 for the N-NTD:dsTRS complex suggested a wide motion between the basic finger and the 2-β5 loop, located at the palm, similar to a tweezer. Interestingly, this tweezer-like motion was intrinsic to the residues located at the dsRNA-binding cleft in N-NTD (Figure 1A).

Our results of conformational flexibility from RMSF and PCA for free and dsRNA-bound N-NTD corroborated each other and suggest a significant contribution of the N- and C-termini and the basic finger (β2-β3 loop) to N-NTD dynamics. They also revealed that N-NTD interaction with dsTRS led to a general gain in protein conformational flexibility when compared to its free state. We suggest that this flexibility gain of dsTRS-bound N-NTD over 25 replicas of concatenated simulations may be a key structural factor to promote dsTRS WC base-pairing destabilization upon N-NTD binding, as determined by the break of intramolecular hydrogen bonds (Figure 2B and 3A) and perturbation of the local base-pair parameters (Figure 4).

### Modeling the dsRNA melting activity

Based on the molecular dynamics simulations performed here, we postulated that only one molecule of N-NTD is necessary to break the WC base-pairing of one molecule of dsRNA, possibly initiating dsRNA melting. To investigate whether this hypothesis is compatible with experimental measurements of N-NTD-induced dsRNA melting, we simulated the experimental data obtained by Grossoehme et al. (2009) (10) using two contrasting kinetic models: (1) model 1, which assumes that melting activity is the result of binding of one N-NTD to one dsRNA; and (2) model 2, which assumes that two N-NTD molecules bind to one dsRNA (sandwich model).

In their work, Grossoehme et al. (2009) measured dsRNA melting activity of N-NTD using fluorescent resonance energy transfer (FRET) from 5’ Cy3-labeled sense RNA strand (TRS) and 3’ Cy5-labeled antisense RNA strand (cTRS) (10). In those experiments, the highest FRET efficiency (∼0.9) was obtained for dsRNA in the absence of N-NTD. Increasing N-NTD concentration led to the dsRNA melting curve, which is characterized by an exponential decay of FRET efficiency as a function of N-NTD concentration. The melting curves reached either zero, for an N-NTD construct that contains the C-terminal serine/arginine (SR)-rich motif, or a plateau, for N-NTD itself (10).

Since the FRET efficiency is a measure of the molar fraction of dsRNA, in the simulated kinetic models presented here, we report the molar fraction of dsRNA as a function of N-NTD concentration, simulating the dsRNA melting curve. We used the software Kinetiscope (http://hinsberg.net/kinetiscope/), which is based on a stochastic algorithm developed by Bunker (24) and Gillespie (25). We used the elementary rate constants for individual chemical steps to produce an absolute time base (Figure 6A). The starting condition mimics exactly the experimental condition, varying the concentration of N-NTD over 50 nM dsRNA (dsTRS). The predictions were validated by direct comparison to the experimental data (10).

**Figure 6.**
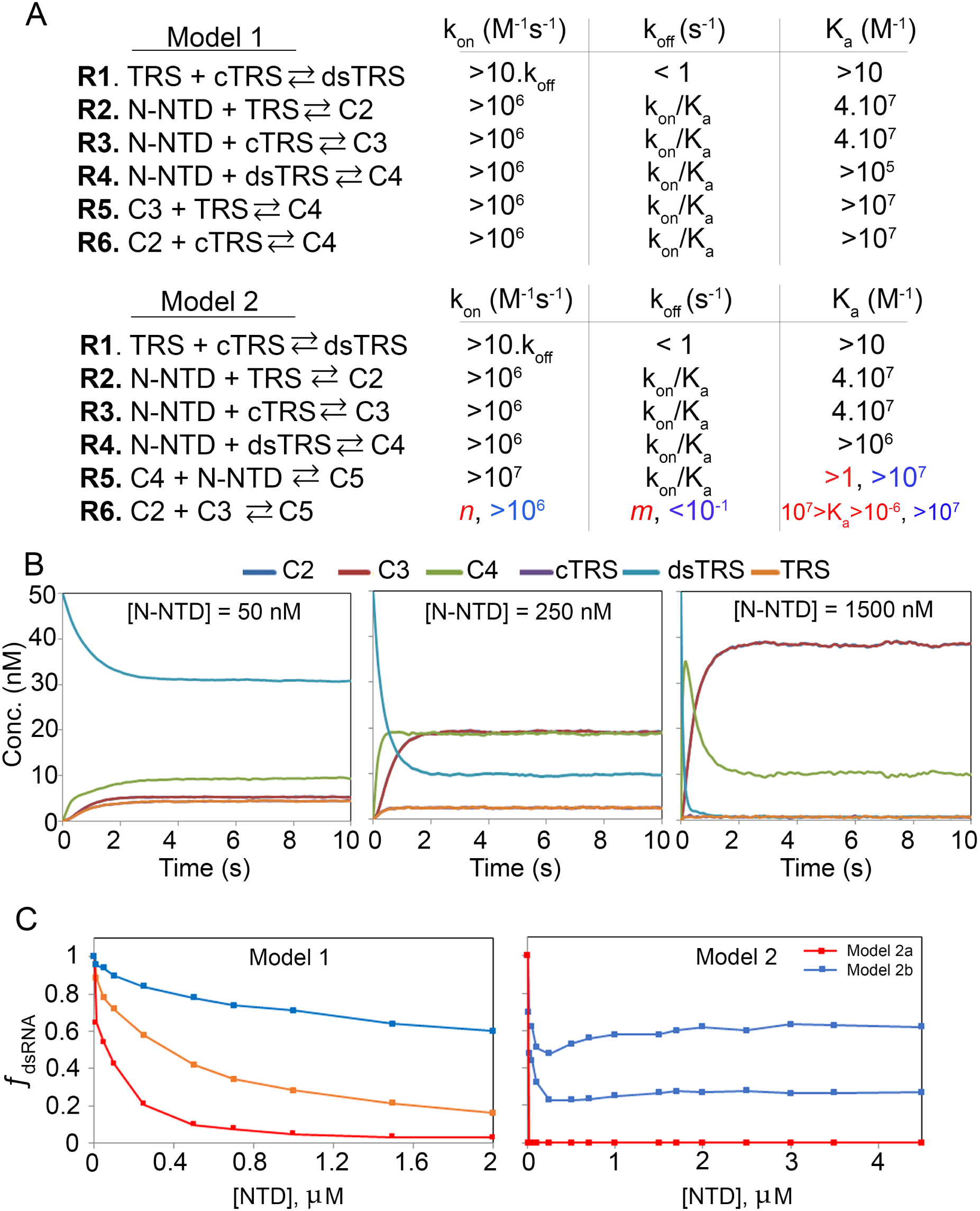
Simulation of the kinetics of dsRNA-melting activity. (A) Reaction R1 to R6 for models 1 and 2. Model 1 implies the melting activity with stoichiometry of 1 N-NTD for 1 dsRNA (C4) and model 2 implies the formation of a sandwich with a stoichiometry two N-NTDs and one dsRNA (C5). At the right of each reaction are the ranges of *k*_*on*_, *k*_*off*_, and *K*_*a*_ in which the simulation produces a dsRNA-melting curve, respecting the boundaries described in the text. For reaction R6 of model 2, the color code refers to the color of the simulated melting curves for model 2. (B) Illustration of the kinetics of dsRNA-melting for model 1 in three different concentrations of N-NTD (50, 250, and 1500 nM). (C) Simulated dsRNA-melting curve for model 1 and 2. We used the starting concentration of 50 nM of dsRNA for all simulations. For model 1 simulations, we used the following reaction rates: *k*_*on*_ (R1) = 4×10^−1^ M^−1^s^−1^ and *k*_*off*_ = 8×10^−4^ s^−1^; *k*_*on*_ (R2, R3) = 4×10^7^ M^−1^s^−1^ and *k*_*off*_ = 1 s^−1^; *k*_*on*_ (R4) = 1×10^7^ M^−1^s^−1^ and *k*_*off*_ = 1 s^−1^; *k*_*on*_ (R5, R6) = 4×10^7^ M^−1^s^−1^ and *k*_*off*_ = 1 s^−1^ (red); *k*_*on*_ (R5, R6) = 4×10^8^ M^−1^s^−1^ and *k*_*off*_ = 1 s^−1^ (orange); *k*_*on*_ (R5, R6) = 6×10^8^ M^−1^s^−1^ and *k*_*off*_ = 1×10^−1^ s^−1^ (blue). For model 2 simulations, we used the following reaction rates: *k*_*on*_ (R1) = 4×10^−1^ M^−1^s^−1^ and *k*_*off*_ = 8×10^−4^ s^−1^; *k*_*on*_ (R2, R3) = 4×10^7^ M^−1^s^−1^ and *k*_*off*_ = 1 s^−1^; *k*_*on*_ (R4) = 1×10^7^ M^−1^s^−1^ and *k*_*off*_ = 1 s^−1^; *k*_*on*_ (R5) = 1×10^8^ M^−1^s^−1^ and *k*_*off*_ = 1 s^−1^ (red); *k*_*on*_ (R6) = 1×10^−1^ M^−1^s^−1^ and *k*_*off*_ = 1×10^8^ s^−1^ (red); *k*_*on*_ (R6) = 1×10^6^ M^−1^s^−1^ and *k*_*off*_ = 1×10^−1^ s^−1^ (blue, bottom); *k*_*on*_ (R6) = 1×10^7^ M^−1^s^−1^ and *k*_*off*_ = 1×10^−1^ s^−1^ (blue, top).

To simulate the melting curve, we had to constrain the kinetic space, which is large because each model is composed by 6 reactions and 12 individual rate constants, assuming the following boundaries: (B1) the kinetic model must be complete, complying all possible reactions for a given mechanism; (B2) the presence of N-NTD must lead to catalysis, with the melting of dsRNA being faster than the annealing reaction; (B3) the equilibrium of the annealing is shifted toward the dsRNA; and (B4) the equilibrium for the melting activity must be reached in few seconds or less to be efficient in the cellular environment.

The criterion for choosing the rate constants for the annealing reaction (R1, Figure 6) was that it must be significantly slower than the melting activity (catalysis). Since to our knowledge, there is no experimental kinetic rate constant available for the annealing of dsTRS, we fixed for the simulations a *k*_*off*_ = 8×10^−4^ s^-1^, which is the experimental value of the dissociation rate constant observed for the almost inactive Y127A N-NTD+SR mutant (10). This mutant has a melting activity of hours, while our simulation showed melting activities of few seconds (Figure 6B). To yield an equilibrium shifted toward the dsRNA, we used *k*_*on*_ = 4×10^−1^ M^−1^s^−1^, which is true below the melting temperature of the dsRNA. Any values of *k*_*off*_ < 1 s^−1^, with an association constant *K*_*a*_, gives the same molar fraction of dsRNA.

We constrained the binding reactions R2 and R3 of N-NTD to the sense (TRS) and antisense (cTRS) single-stranded RNA (ssRNA) (Figure 6A) based on the published experimental values for these association constants (10, 21). Since these values were very similar, to simplify the simulation, we used the same *K*_*a*_ for both reactions (*K*_*a*_ = 4×10^7^ M^− 1^). Note that *k*_*on*_ < 10^6^ M^-1^s^-1^ makes the reaction too slow to reach equilibrium, violating boundary B4 (Figures S14A and S16B).

For dsRNA (dsTRS) binding, there was no experimental data to constrain the simulation. However, simulations unambiguously showed that *K*_*a*_ for reaction R4 must be of the same order of that for ssRNAs, leading to the allowed ranges depicted in Figure 6A. We also determined *k*_*on*_ based on the simulations, taking boundaries B2 and B4 into consideration, which were also considered for reactions R1 to R3 (Figure S14B, S15A and S16B).

All the constraints applied to reactions R1 to R4 are valid for both kinetic models (models 1 and 2). Conversely, reactions R5 and R6 are specific for each kinetic model, being essential to comply with boundary B1.

For model 1, there is no experimental data available to constrain reactions R5 and R6, but the simulations showed that they are tightly related to reactions R2 and R3, being both *K*_*a*_ and *k*_*off*_ of the same order of magnitude for reactions R2 and R3 (Figure S14C). Note that there is an intricate relationship between the formation of ssRNA-bound states (C2 and C3) and the decrease of free or bound dsRNA (dsTRS and C4). To illustrate this relationship, Figure 6B shows the kinetics at three concentrations of N-NTD. The simulated melting curves for model 1 resembled the near exponential decay observed experimentally (Figure 6C, left). Interestingly, when *k*_*off*_ of reactions R5 and R6 were bigger than *k*_*off*_ for reactions R2 and R3, we observed a plateau in the exponential decay of the dsRNA melting curve (Figure 6C). Remarkably, melting curves that either decayed to zero or reached a plateau was observed experimentally, as mentioned before (10). It is worth mentioning that the kinetic model 1 is fully compatible with the experimental data by Grossoehme et al. (2009) (10), as well as with the mechanism suggested by the MD simulations, in which one N-NTD can initiate dsRNA melting, destabilizing the WC base-pairing.

We also evaluated kinetic model 2. This mechanism for N-NTD melting activity was suggested in the conclusion scheme drawn in the paper by Grossoehme et al. (2009) (10), in which each N-NTD of the dimeric full-length nucleocapsid protein binds to one dsRNA to catalyze the melting reaction. In this model, a sandwich of two N-NTDs and one dsRNA is formed, and the final products are each N-NTD bound to TRS and cTRS ssRNA.

To build a kinetic model that would exclusively produce ssRNA from the sandwiched dsRNA, we had to replace reactions R5 and R6 of kinetic model 1. In the new model, reaction R5 forms the sandwiched dsRNA (C5, Figure 6A) and reaction R6 is the dissociation of C5 into the ssRNA-bound N-NTDs (C2 and C3, Figure 6A). To simulate N-NTD melting activity considering model 2, we used the same boundaries described earlier (B1, B2, B3 and B4), with reactions R1 to R4 having almost the same constraints described for model 1.

Reaction R5 and R6 of model 2 has no parallel to any other reaction. We scanned all the kinetic space that led to the catalysis of melting activity and observed two contrasting situations. The first is when reaction R6 equilibrium is between 10^−6^ and 10^7^ M^-1^, always having the dissociated forms C2 and C3 available and making the melting curve very stiff (model 2a). The second is the opposite situation, where equilibrium is skilled toward the sandwich state (C5) with *K*_*a*_ > 10^7^ M^−1^ (model 2b). Figure 6C illustrates the melting curves obtained for the two situations.

Model 2a is characterized for the high efficiency in the dissociation of the dsRNA, *k*_*on*_ and *k*_*off*_ can assume any value. Particularly for model 2a, the kinetic of dsRNA melting is also independent of *k*_*on*_ for reactions R2 and R3, at fixed concentrations of N-NTD. All simulated conditions led to the curve in red (Figure 6C), in which, the minimal amount of N-NTD (10 nM) led to complete dissociation of the dsRNA (molar fraction of zero). Figure S15 illustrates all the simulated boundaries. Note that for model 2a, there is never an accumulation of C5 (Figure S15C).

Model 2b corresponds to when the equilibrium of reaction R6 is shifted toward C5 (*K*_*a*_ > 10^7^ M^−1^). Figure S16 illustrates the reaction boundaries. In this situation, we were able to observe a melting curve (Figure 6C, blue) with a near exponential decay at a low concentration of N-NTD and a near exponential rise at higher concentrations of N-NTD. This behavior is explained by the accumulation of C5 and N-NTD concentration-dependent mutual compensation of C5 and dsRNA. Note how *K*_*a*_ modulates the accumulation of C5, transitioning between models 2a and 2b (Figure S16D). The increase in the concentration of N-NTD led to a decrease in dsRNA forming C2 and C3 and ssRNA. Further increase in N-NTD led to a decrease in dsRNA and a compensating increase in C5. None of the situations simulated for model 2 are parallel to the experimental observation.

Reaction R5 of model 2, which forms the sandwiched dsRNA, follows reaction R4 for model 2b. We determined that *k*_*on*_ has to be > 10^7^ s^−1^ to keep up with boundaries B2 and B4. For the melting activity to take place, the equilibrium of reaction R5 was shifted toward C5 (*K*_*a*_ > 1, for model 2a or 10^7^ M^−1^ for model 2b) (Figure S15B and S16C).

## DISCUSSION

In the present work, we used computational simulations to unravel the dsRNA melting activity of the isolated SARS-CoV-2 N-NTD. Our molecular dynamics data suggested that, during interaction with dsRNA, protein dynamics drives the destabilization of hydrogen bonds involved in the WC RNA base-pairing, probably in a 1:1 stoichiometry (N-NTD:dsRNA). We also showed that the capacity of N-NTD to break the WC base-pairing was sequence-specific, being more efficient for dsTRS (5’–UCUAAAC–3’) than for a non-specific (NS) sequence (5’–CACUGAC–3’). To further explore the N-NTD:dsRNA stoichiometry, we constructed kinectic models based on experimental data by Grossoehme et al (2009) (10). Remarkably, the model using a 1:1 stoichiometry greatly fits the experimental data, reinforcing the mechanism we hypothesize here.

The strategy of performing 25 100 ns-molecular dynamics simulations with the same starting structure but different seeds of the random number generator provided a large sampling of conformational space of each molecular system (N-NTD, dsRNAs, and N-NTD/dsRNA complexes). This set of theoretical data ensured a statistically significant result showing that N-NTD destabilizes the WC base-pairing, especially for dsTRS, through the replacement of intramolecular hydrogen bonds between the dsRNA strands by intermolecular hydrogen bonds between N-NTD and the nitrogenous bases of each RNA strand. The results also revealed unbiasedly that the rigid-body geometric parameters of the WC base-pairing were significantly changed due to N-NTD binding.

One notable N-NTD structural feature is the presence of a significant number of loops; only 32 out of 140 residues are involved in secondary structure (16, 26). This is a typical feature of a dynamic protein. In fact, our results revealed that N-NTD is a plastic protein, with the N and C-termini and the β2-β3 loop (finger) as the most prominent dynamic regions. For the N-NTD:dsTRS interaction, a remarkable tweezer-like motion between the finger and the 2-β5 loop could be related to the sequence-specific WC base-pairing destabilization. This led us to hypothesize that, following formation of the N-NTD:dsTRS complex, the tweezer-like motion resulted from intrinsic protein dynamics might promote a steric effect causing a “compaction pressure” on the dsRNA strands. This might expose residues from the bottom of palm (finger/ 2-β5 cleft) allowing their interaction with the bases, leading to destabilization of the WC base-pairing (Figure 7).

**Figure 7.**
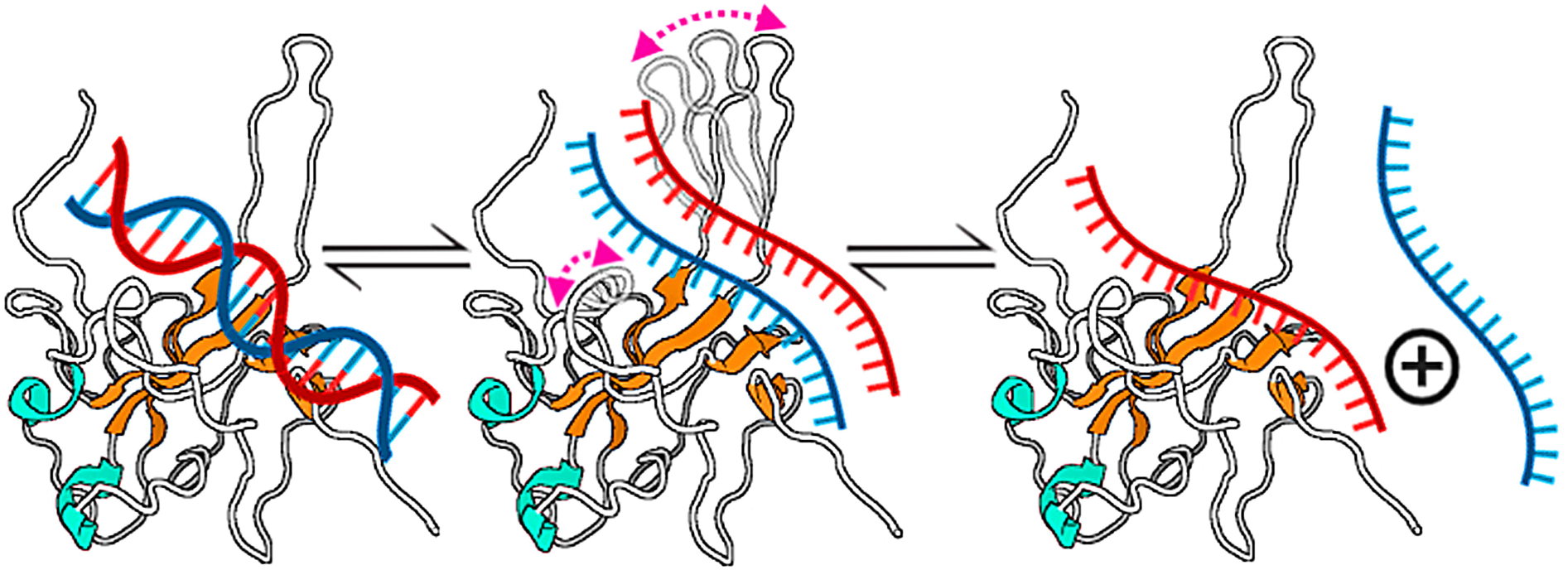
Summary of the proposed mechanism for dsRNA-melting activity of N-NTD. The binding of one N-NTD to one dsRNA triggers the destabilization of WC base-pairing of the dsRNA and consequently expose the nitrogenous bases for interacting directly with the N-NTD. We suggest that this activity is a consequence of intrinsic dynamics of N-NTD, especially because the tweezer-like motion between β2-β3 (finger) and 2-β5 loops. The protein is denoted as cartoon with the helix- and β-strand secondary structures colored in cyan and orange, respectively. The dsRNA is showed as a line model with the complementary strands colored in red and blue. The tweezer-like motion between the finger and 2-β5 loop is indicated by bidirectional arrows colored in magenta.

To confirm the mechanism emerged from the MD simulations, where only one molecule of N-NTD was enough to initiate dsRNA melting, we constructed kinetic models considering two possible scenarios: a stoichiometry of 1:1 or 2:1 for N-NTD and dsRNA. The 2:1 stoichiometry is more intuitive since N protein is dimeric in solution (27). However, the 1:1 stoichiometry produced dsRNA-melting curves compatible with the experimental data published by Grossoehme et al. (2009) (10). It is important to mention that the simulations of the kinetic models were only possible by constraining the kinetic space since the number of degrees of freedom for 6 reactions is quite large. This constrained kinetic space was created by imposing boundaries to the system. These simulations brought two important conclusions: (i) the 1:1 stoichiometry is enough to explain the experimental data; and (ii) the sandwich model 2 is less likely to occur since simulations produced more complex melting curves, which are different from the experimental data.

Altogether, the results presented here support the idea that two N-NTDs of dimeric N protein would not be necessary to act on one dsRNA motif (dsTRS). Each N-NTD of the same dimer would work independently, leading to a gain in efficiency of full-length N when compared to the sandwich model (model 2). The observation that one N-NTD could be able to melt dsRNA opens two new avenues for the understanding of the role played by the N protein in the viral replication cycle. First, the two N-NTD arms of full-length N could bind to two different sites at the viral genome, making it possible to bridge two distant regions (2:2 stoichiometry). Second, if monomeric full-length protein is active, a regulatory event involving the dissociation of the N protein dimer would be consider. Indeed, for the N protein of bovine betacoronavirus, Lo et al. (2017) suggested that it acts as a bridge between distant motifs in the genome.

## MATERIAL AND METHODS

### Molecular docking

To perform the docking, we took advantage of experimental data previously published by Dinash et al. (2020), in which SARS-CoV-2 N-NTD interaction with a non-specific (NS) dsRNA (5’–CACUGAC–3’) was monitored by chemical shift perturbations (CSPs) titration experiments (16). Structural models for N-NTD:dsNS complex were constructed using the HADDOCK server (version 2.2) (20). The coordinates used as input were obtained from the solution NMR structure of SARS-CoV-2 N-NTD (PDB 6YI3) (16), and the X-ray structure of a synthetic 7-mer dsRNA (PDB 4U37) (28) mutated using the w3DNA server (version 2.0) (29) to generate the same NS dsRNA sequence as that used in CSP titration experiments (5’–CACUGAC–3’). In addition, histidine protonation states were set according to the PROPKA server (30) considering pH 7.0. In total, 2000 complex structures of rigid-body docking were calculated by using the standard HADDOCK protocol with an optimized potential for liquid simulation (OPLSX) parameters (31). The final 200 lowest-energy structures were selected for subsequent explicit solvent (water) and semi-flexible simulated annealing refinement, to optimize side chain constants. The N-NTD:dsNS complex we obtained is very similar to that reported by Dinash et al. (2020) (16).

Next, the structural model of the N-NTD:dsTRS (5’–UCUAAAC–3’) complex was generated from the lowest-energy structure of the N-NTD:dsNS complex, derived from the cluster with the lowest HADDOCK score, by mutating the dsRNA sequence using w3DNA (29). Therefore, both complexes have identical geometries, varying only the dsRNA sequences. Structural conformation of the constructed model for N-NTD:dsTRS complex was displayed using the web application http://skmatic.x3dna.org for easy creation of DSSR (Dissecting the Spatial Structure of RNA)-PyMOL schematics (32).

### Molecular dynamics simulation

Molecular dynamics (MD) calculations for N-NTD, dsRNAs, and N-NTD:dsRNA complexes were performed using GROMACS (version 5.0.7) (33). The molecular systems were modeled with the corrected AMBER14-OL15 package, including the ff14sb protein (34) and ff99bsc0χ_OL3_+OL15 nucleic acid (35, 36) force fields, as well as the TIP3P water model (37). The structural models of N-NTD (PDB 6YI3), dsRNAs (mutated PDB 4U37), and N-NTD:dsRNA complexes (from molecular docking) were placed in the center of a cubic box solvated by a solution of 50 mM NaCl in water. The protonation state of ionizable residues was set according to the PROPKA server (30) considering pH 7.0. Periodic boundary conditions were used and all simulations were performed in NPT ensemble, keeping the system at 298 K and 1.0 bar using Nose-Hoover thermostat (*τ*_*T*_ = 2 ps) and Parrinello-Rahman barostat (*τ*_*P*_ = 2 ps and compressibility = 4.5×10^−5^·bar^−1^). A cutoff of 12 Å for both Lennard-Jones and Coulomb potentials was used. The long-range electrostatic interactions were calculated using the particle mesh Ewald (PME) algorithm. In every MD simulation, a time step of 2.0 fs was used and all covalent bonds involving hydrogen atoms were constrained to their equilibrium distance. A conjugate gradient minimization algorithm was used to relax the superposition of atoms generated in the box construction process. Energy minimizations were carried out with steepest descent integrator and conjugate gradient algorithm, using 1,000 kJ·mol^−1^·nm^−1^ as maximum force criterion. One hundred thousand steps of molecular dynamics were performed for each NVT and NPT equilibration, applying force constants of 1,000 kJ·mol^−1^·nm^−2^ to all heavy atoms of N-NTD, dsRNAs, and N-NTD:dsRNA complexes. At the end of preparation, 25 replicas of 100 ns MD simulation of each molecular system with different seeds of the random number generator were carried out for data acquisition, totalizing 2.5 μs. Following dynamics, the trajectories of each molecular system were firstly concatenated individually and analyzed according to the root mean square deviation (RMSD) of backbone atoms of protein and nucleic acid, the number of contacts (< 0.6 nm) between N-NTD and dsRNAs, the intramolecular (dsNS and dsTRS strands) and intermolecular (N-NTD:dsRNA complexes) hydrogen bonds (cutoff distance = 3.5 Å and maximum angle = 30°), and the local base-pair parameters [angles (Å): buckle, opening, and propeller; distances (nm): stretch, stagger, and shear] for free and N-NTD-bound dsNS and dsTRS by using the *do_x3dna* tool (22) along with the 3DNA package (23). These local base-pair parameters of each of the 25 runs and of selected replicas were analyzed together as histogram plots exhibiting population distributions. After individual analysis of each simulation, the last 50 ns of the 25 trajectories of free and dsRNA-bound N-NTD were concatenated in single files, and these new trajectories were used to evaluate the root mean square fluctuation (RMSF) of the C atoms and principal component analysis (PCA). PCA scatter plots were generated for free and dsRNA-bound N-NTD, as well as conformational motions were filtered (30 frames) from the eigenvectors of the first and second principal components (PC1 and PC2, respectively). The structural representations of the motions from PC1 and PC2 were prepared using PyMol (38).

### Kinetic simulations

The Kinetiscope program (version 1.1.956.x6, http://hinsberg.net/kinetiscope/) was used to simulate the kinetics of dsRNA melting by SARS-CoV-2 N-NTD. Simulations were performed under constant volume, pressure, and temperature (298.15 K). An initial concentration of 50 nM dsTRS was used and a total of 2439 initial number of particles were calculated. N-NTD concentration ranged from 0 to 2 μM for model 1 and from 0 to 4.5 μM for model 2. The maximum number of events was set to ten million and simulations lasted 100 s.

## ACKNOWLEDGMENTS

The author I.P.C. gratefully acknowledges the financial support by postdoctoral fellowship from FAPERJ and the PROPe UNESP. The authors are grateful for the access to the Santos Dumont supercomputer at the National Laboratory of Scientific Computing (LNCC), Brazil.

## REFERENCES

1. Zhu N, Zhang D, Wang W, Li X, Yang B, Song J, Zhao X, Huang B, Shi W, Lu R, Niu P, Zhan F, Ma X, Wang D, Xu W, Wu G, Gao GF, Tan W. 2020. A novel coronavirus from patients with pneumonia in China, 2019. N Engl J Med 382:727–733.

2. Zhou P, Yang X Lou, Wang XG, Hu B, Zhang L, Zhang W, Si HR, Zhu Y, Li B, Huang CL, Chen HD, Chen J, Luo Y, Guo H, Jiang R Di, Liu MQ, Chen Y, Shen XR, Wang X, Zheng XS, Zhao K, Chen QJ, Deng F, Liu LL, Yan B, Zhan FX, Wang YY, Xiao GF, Shi ZL. 2020. A pneumonia outbreak associated with a new coronavirus of probable bat origin. Nature 579:270–273.

3. Klein S, Cortese M, Winter SL, Wachsmuth-melm M, Neufeldt CJ, Stanifer ML, Steeve B, Bartenschlager R, Chlanda P. 2020. SARS-CoV-2 structure and replication characterized by in situ cryo-electron tomography. BioRxiv 10.1101/20:1-8.

4. Naqvi AAT, Fatima K, Mohammad T, Fatima U, Singh IK, Singh A, Atif SM, Hariprasad G, Hasan GM, Hassan MI. 2020. Insights into SARS-CoV-2 genome, structure, evolution, pathogenesis and therapies: Structural genomics approach. Biochim Biophys Acta - Mol Basis Dis 1866:165878.

5. Fang SG, Shen H, Wang J, Tay FPL, Liu DX. 2008. Proteolytic processing of polyproteins 1a and 1ab between non-structural proteins 10 and 11/12 of Coronavirus infectious bronchitis virus is dispensable for viral replication in cultured cells. Virology 379:175–180.

6. Snijder EJ, van der Meer Y, Zevenhoven-Dobbe J, Onderwater JJM, van der Meulen J, Koerten HK, Mommaas AM. 2006. Ultrastructure and Origin of Membrane Vesicles Associated with the Severe Acute Respiratory Syndrome Coronavirus Replication Complex. J Virol 80:5927–5940.

7. Snijder EJ, Bredenbeek PJ, Dobbe JC, Thiel V, Ziebuhr J, Poon LLM, Guan Y, Rozanov M, Spaan WJM, Gorbalenya AE. 2003. Unique and conserved features of genome and proteome of SARS-coronavirus, an early split-off from the coronavirus group 2 lineage. J Mol Biol 331:991–1004.

8. Thiel V, Ivanov KA, Putics Á, Hertzig T, Schelle B, Bayer S, Weißbrich B, Snijder EJ, Rabenau H, Doerr HW, Gorbalenya AE, Ziebuhr J. 2003. Mechanisms and enzymes involved in SARS coronavirus genome expression. J Gen Virol. J Gen Virol.

9. Kuo L, Hurst-Hess KR, Koetzner CA, Masters PS. 2016. Analyses of Coronavirus Assembly Interactions with Interspecies Membrane and Nucleocapsid Protein Chimeras. J Virol 90:4357–4368.

10. Grossoehme NE, Li L, Keane SC, Liu P, Dann CE, Leibowitz JL, Giedroc DP. 2009. Coronavirus N Protein N-Terminal Domain (NTD) Specifically Binds the Transcriptional Regulatory Sequence (TRS) and Melts TRS-cTRS RNA Duplexes. J Mol Biol 394:544–557.

11. Verheije MH, Hagemeijer MC, Ulasli M, Reggiori F, Rottier PJM, Masters PS, Haan CAM de. 2010. The Coronavirus Nucleocapsid Protein Is Dynamically Associated with the Replication-Transcription Complexes. J Virol 84:11575–11579.

12. Zúñiga S, Cruz JLG, Sola I, Mateos-Gómez PA, Palacio L, Enjuanes L. 2010. Coronavirus Nucleocapsid Protein Facilitates Template Switching and Is Required for Efficient Transcription. J Virol 84:2169–2175.

13. Chang C-K, Hsu Y-L, Chang Y-H, Chao F-A, Wu M-C, Huang Y-S, Hu C-K, Huang T-H. 2009. Multiple Nucleic Acid Binding Sites and Intrinsic Disorder of Severe Acute Respiratory Syndrome Coronavirus Nucleocapsid Protein: Implications for Ribonucleocapsid Protein Packaging. J Virol 83:2255–2264.

14. Surjit M, Liu B, Kumar P, Chow VTK, Lal SK. 2004. The nucleocapsid protein of the SARS coronavirus is capable of self-association through a C-terminal 209 amino acid interaction domain. Biochem Biophys Res Commun 317:1030–1036.

15. McBride R, van Zyl M, Fielding BC. 2014. The coronavirus nucleocapsid is a multifunctional protein. Viruses 6:2991–3018.

16. Dinesh DC, Chalupska D, Silhan J, Veverka V, Boura E. 2020. Structural basis of RNA recognition by the SARS-CoV-2 nucleocapsid phosphoprotein. bioRxiv 2020.04.02.022194.

17. Sola I, Almazán F, Zúñiga S, Enjuanes L. 2015. Continuous and Discontinuous RNA Synthesis in Coronaviruses. Annu Rev Virol 2:265–288.

18. Zúñiga S, Sola I, Alonso S, Enjuanes L. 2004. Sequence Motifs Involved in the Regulation of Discontinuous Coronavirus Subgenomic RNA Synthesis. J Virol 78:980–994.

19. Chang CK, Hou MH, Chang CF, Hsiao CD, Huang TH. 2014. The SARS coronavirus nucleocapsid protein - Forms and functions. Antiviral Res. Elsevier.

20. Van Zundert GCP, Rodrigues JPGLM, Trellet M, Schmitz C, Kastritis PL, Karaca E, Melquiond ASJ, Van Dijk M, De Vries SJ, Bonvin AMJJ. 2016. The HADDOCK2.2 Web Server: User-Friendly Integrative Modeling of Biomolecular Complexes. J Mol Biol 428:720–725.

21. Keane SC, Lius P, Leibowitzs JL, Giedroc DP. 2012. Functional Transcriptional Regulatory Sequence (TRS) RNA binding and helix destabilizing determinants of Murine Hepatitis Virus (MHV) Nucleocapsid (N) protein. J Biol Chem 287:7063–7073.

22. Kumar R, Grubmüller H. 2015. Structural bioinformatics do_x3dna: a tool to analyze structural fluctuations of dsDNA or dsRNA from molecular dynamics simulations. Bioinformatics 31:2583–2585.

23. Lu XJ, Olson WK. 2008. 3DNA: A versatile, integrated software system for the analysis, rebuilding and visualization of three-dimensional nucleic-acid structures. Nat Protoc 3:1213–1227.

24. Bunker DL, Garrett B, Kleindienst T, Long GS. 1974. Discrete simulation methods in combustion kinetics. Combust Flame 23:373–379.

25. Gillespie DT. 1976. A general method for numerically simulating the stochastic time evolution of coupled chemical reactions. J Comput Phys 22:403–434.

26. Kang S, Yang M, Hong Z, Zhang L, Huang Z, Chen X, He S, Zhou Z, Zhou Z, Chen Q, Yan Y, Zhang C, Shan H, Chen S. 2020. Crystal structure of SARS-CoV-2 nucleocapsid protein RNA binding domain reveals potential unique drug targeting sites. Acta Pharm Sin B.

27. Zeng W, Liu G, Ma H, Zhao D, Yang Y, Liu M, Mohammed A, Zhao C, Yang Y, Xie J, Ding C, Ma X, Weng J, Gao Y, He H, Jin T. 2020. Biochemical characterization of SARS-CoV-2 nucleocapsid protein. Biochem Biophys Res Commun 527:618–623.

28. Sheng J, Larsen A, Heuberger BD, Blain JC, Szostak JW. 2014. Crystal structure studies of RNA duplexes containing s2U:A and s2U:U base Pairs. J Am Chem Soc 136:13916–13924.

29. Li S, Olson WK, Lu X-J. 2019. Web 3DNA 2.0 for the analysis, visualization, and modeling of 3D nucleic acid structures. Web Serv issue Publ online 47.

30. Olsson MHM, SØndergaard CR, Rostkowski M, Jensen JH. 2011. PROPKA3: Consistent treatment of internal and surface residues in empirical p K a predictions. J Chem Theory Comput 7:525–537.

31. Linge JP, Williams MA, Spronk CAEM, Bonvin AMJJ, Nilges M. 2003. Refinement of protein structures in explicit solvent. Proteins Struct Funct Genet 50:496–506.

32. Lu X-J. 2020. DSSR-enabled innovative schematics of 3D nucleic acid structures with PyMOL - PubMed. Nucleic Acids Res 48:e74.

33. Abraham MJ, Murtola T, Schulz R, Páll S, Smith JC, Hess B, Lindah E. 2015. Gromacs: High performance molecular simulations through multi-level parallelism from laptops to supercomputers. SoftwareX 1–2:19–25.

34. Maier JA, Martinez C, Kasavajhala K, Wickstrom L, Hauser KE, Simmerling C. 2015. ff14SB: Improving the Accuracy of Protein Side Chain and Backbone Parameters from ff99SB. J Chem Theory Comput 11:3696–3713.

35. Zgarbová M, Šponer J, Otyepka M, Cheatham TE, Galindo-Murillo R, Jurečka P. 2015. Refinement of the Sugar-Phosphate Backbone Torsion Beta for AMBER Force Fields Improves the Description of Z-and B-DNA. J Chem Theory Comput 11:5723–5736.

36. Zgarbová M, Otyepka M, Šponer J, Mládek A, BanáŠ P, Cheatham TE, Jurečka P. 2011. Refinement of the Cornell et al. Nucleic acids force field based on reference quantum chemical calculations of glycosidic torsion profiles. J Chem Theory Comput 7:2886–2902.

37. Jorgensen WL, Chandrasekhar J, Madura JD, Impey RW, Klein ML. 1983. Comparison of simple potential functions for simulating liquid water. J Chem Phys 79:926–935.

38. Delano WL. 2002. The PyMOL Molecular Graphics System. DeLano Scientific, San Carlos, CA, USA. San Carlos, CA, USA.

39. Lu X-J, Olson WK. 2003. 3DNA: a software package for the analysis, rebuilding and visualization of three-dimensional nucleic acid structures. Nucleic Acids Res 31:5108–5121.

