## Supplementary material for "Dynamics of the N-terminal domain of SARS-CoV-2 nucleocapsid protein drives dsRNA melting in a counterintuitive tweezer-like mechanism"

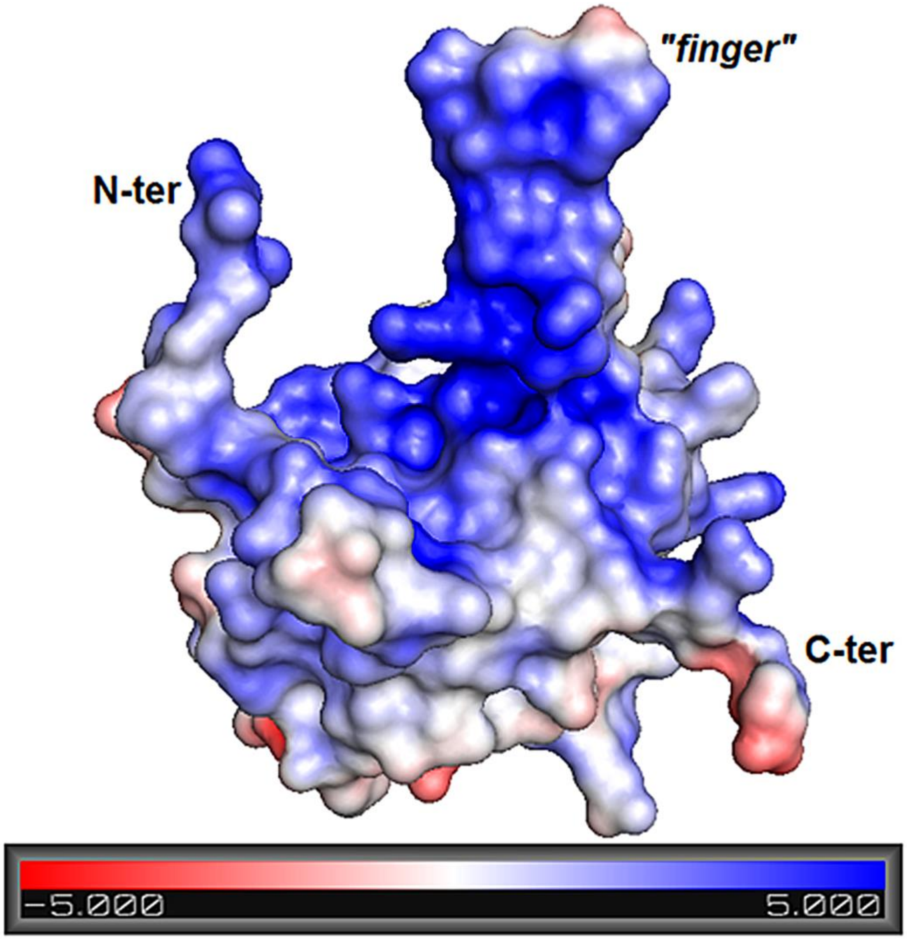


**Figure S1.** Electrostatic potential surface of SARS-CoV-2 N-NTD calculated from APBS software (1) using charge values and protonation states determined by PDB2PQR server (2) along with PROPKA program (pH 7.0, 50 mM NaCl, 25 °C) (3). The bar denotes the electrostatic potential range from –5 (red) to +5 kT (blue). The electrostatic potential surface of N-NTD was displayed using PyMOL (4).

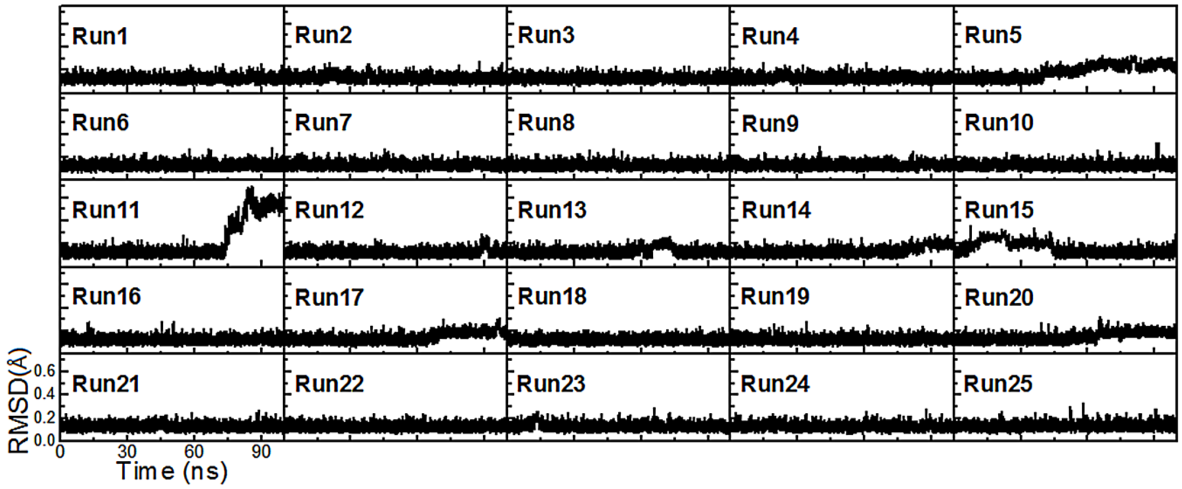


**Figure S2.** RMSD values of the backbone atoms of free dsTRS for the 25 replicas of 100 ns MD simulations.

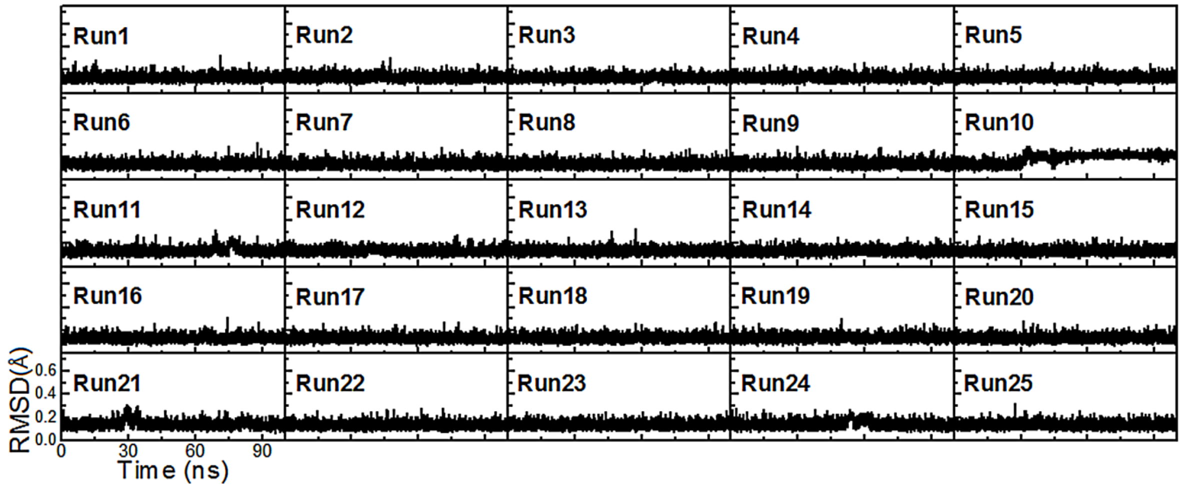


**Figure S3.** RMSD values of the backbone atoms of free dsNS for the 25 replicas of 100 ns MD simulations.

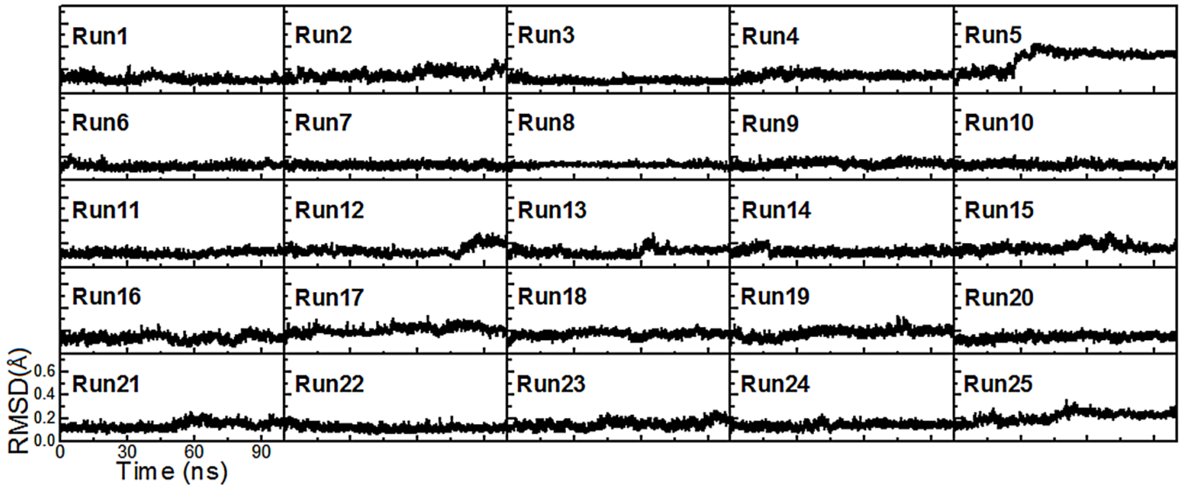


**Figure S4.** RMSD values of the backbone atoms of N-NTD-bound dsTRS for the 25 replicas of 100 ns MD simulations.

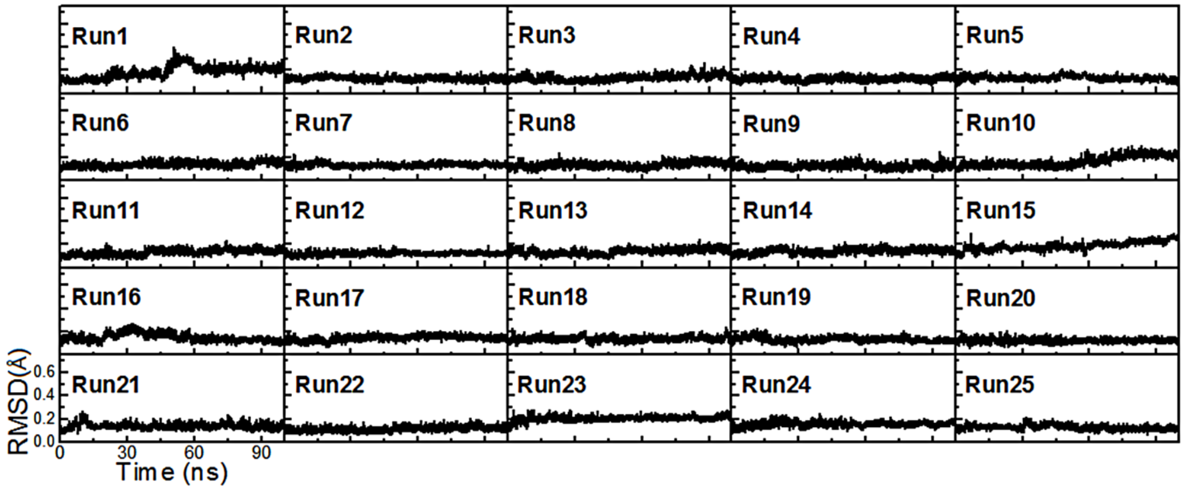


**Figure S5.** RMSD values of the backbone atoms of N-NTD-bound dsNS for the 25 replicas of 100 ns MD simulations.

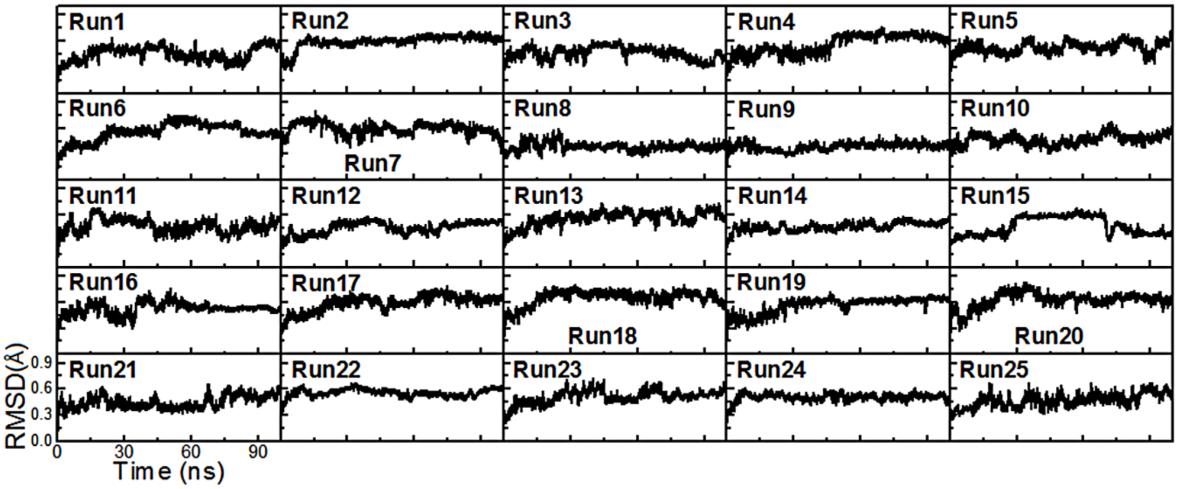


**Figure S6.** RMSD values of the backbone atoms of free N-NTD for the 25 replicas of 100 ns MD simulations.

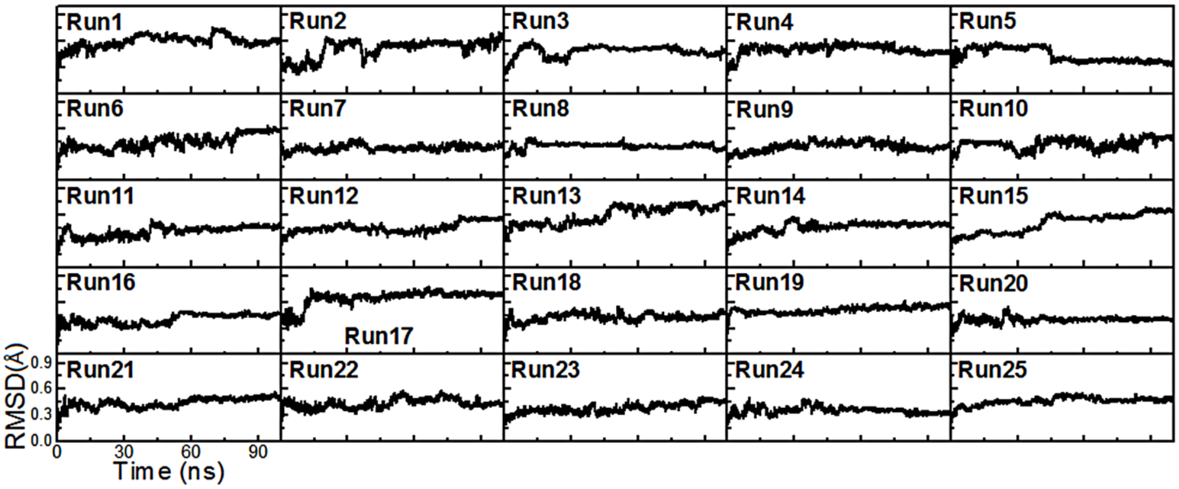


**Figure S7.** RMSD values of the backbone atoms of dsTRS-bound N-NTD for the 25 replicas of 100 ns MD simulations.

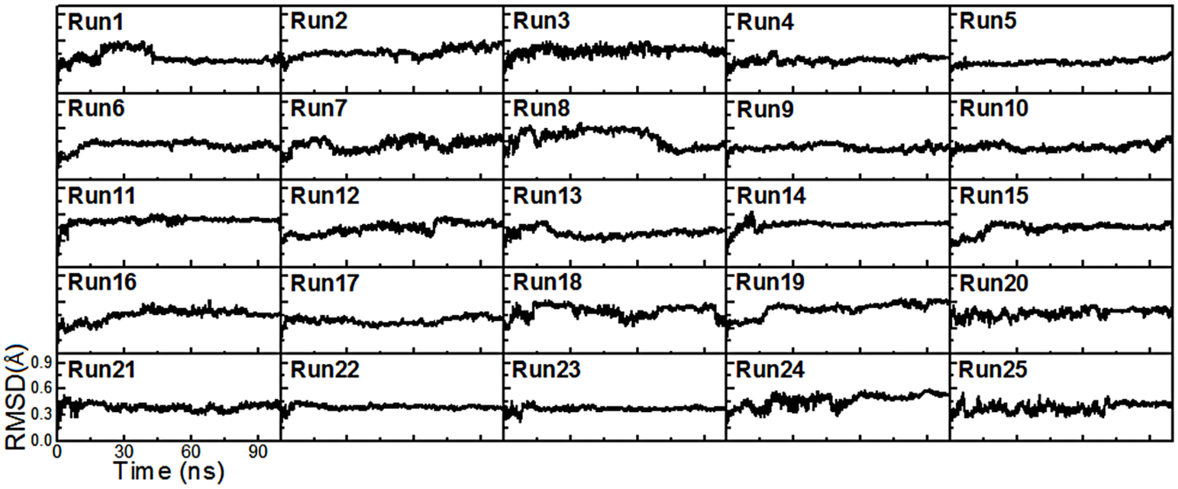


**Figure S8.** RMSD values of the backbone atoms of dsNS-bound N-NTD for the 25 replicas of 100 ns MD simulations.

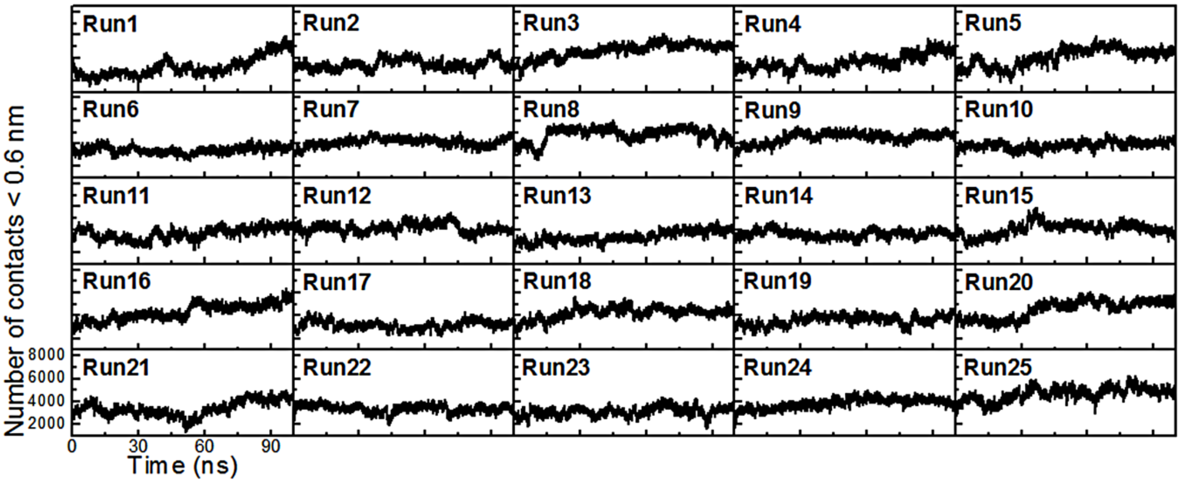


**Figure S9.** Number of contacts < 0.6 nm between the atoms of N-NTD and dsTRS for the 25 replicas of 100 ns MD simulations.

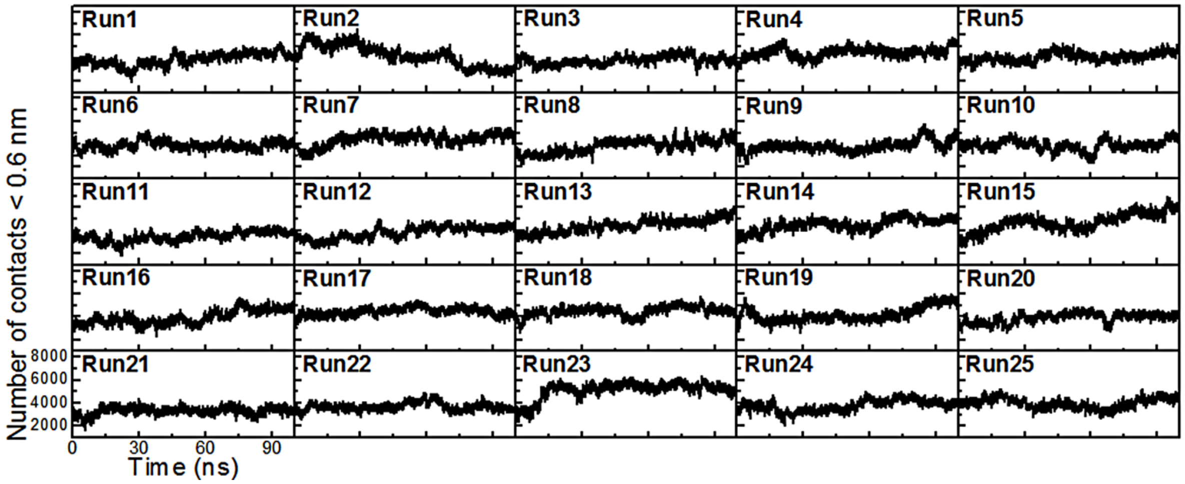


**Figure S10.** Number of contacts < 0.6 nm between the atoms of N-NTD and dsNS for the 25 replicas of 100 ns MD simulations.

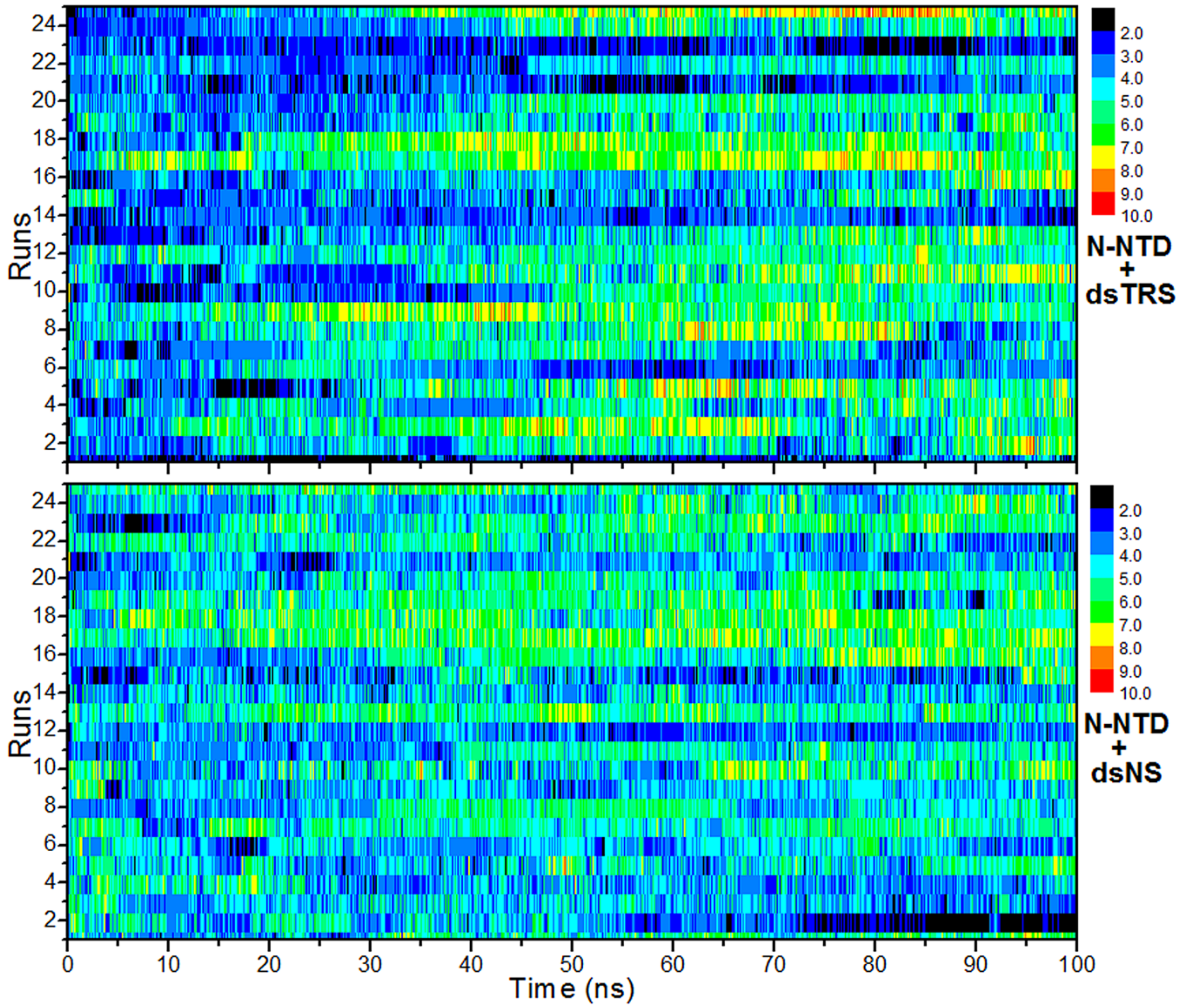


**Figure S11.** Number of intermolecular hydrogen bonds formed between the nitrogenous bases of the dsRNAs (dsTRS in top and dsNS in bottom) and N-NTD over the 100 ns simulations for the 25 MD replicas. The color bar denotes the correspondence between the color code and the number of intermolecular hydrogen bonds.

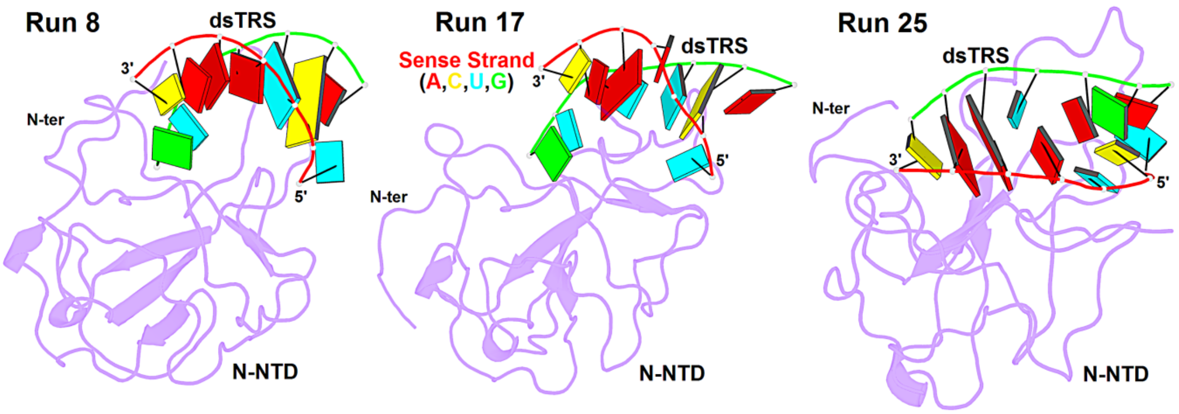


**Figure S12.** Structural model of the N-NTD/dsTRS complex representative of the MD simulation for the runs 8, 17, and 25. The protein is shown as purple cartoon and dsTRS is denoted as ribbon model with nitrogenous bases and base-pairing as colored squares and rectangles, respectively. The color of the squares corresponds to the type of nitrogenous base, being A: red, C: yellow, U: cyan, and G: green, while for the rectangles refer to the nitrogenous base color of the sense strand of dsRNA.

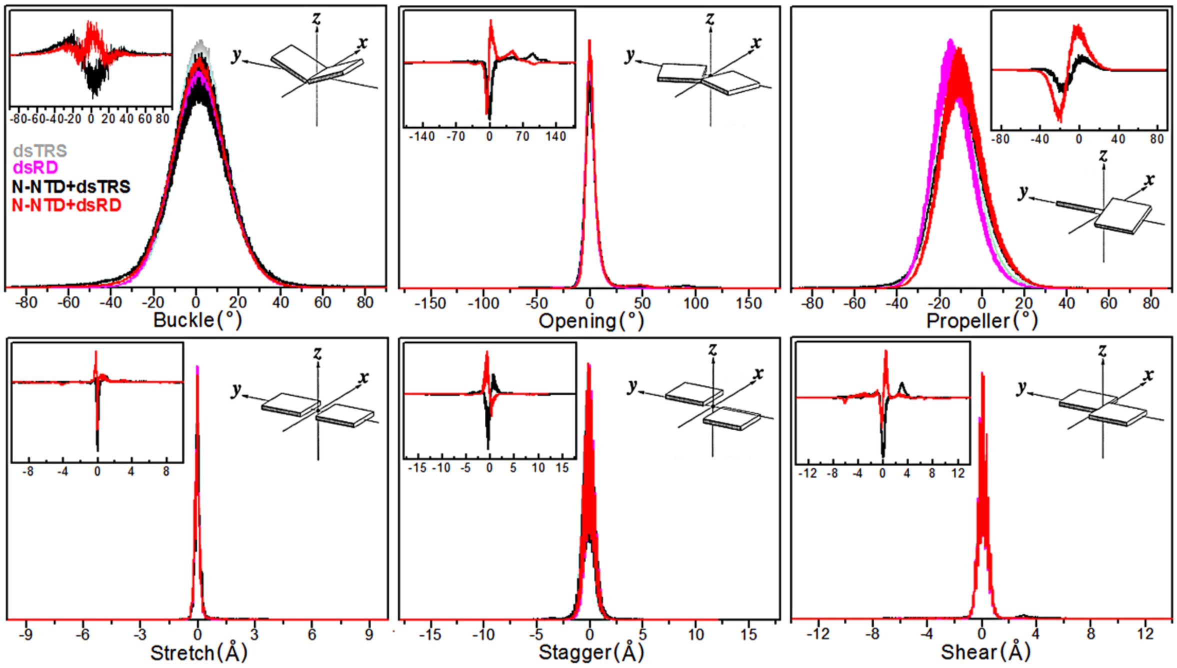


**Figure S13.** Population distributions of local base-pair parameters (angles: buckle, opening, and propeller; distances: stretch, stagger, and shear) for 25 runs of dsTRS and dsNS in their free form (dsTRS in light gray and dsNS in magenta, respectively) and complexed with N-NTD (N-NTD+dsTRS in black and N-NTD+dsNS in red). The plot insets correspond to the difference between the population distributions of N-NTD-bound dsRNA minus its free state for dsNS (red) and dsTRS (black). The scheme insets illustrate the geometrical definition of each local base-pair parameter (5).

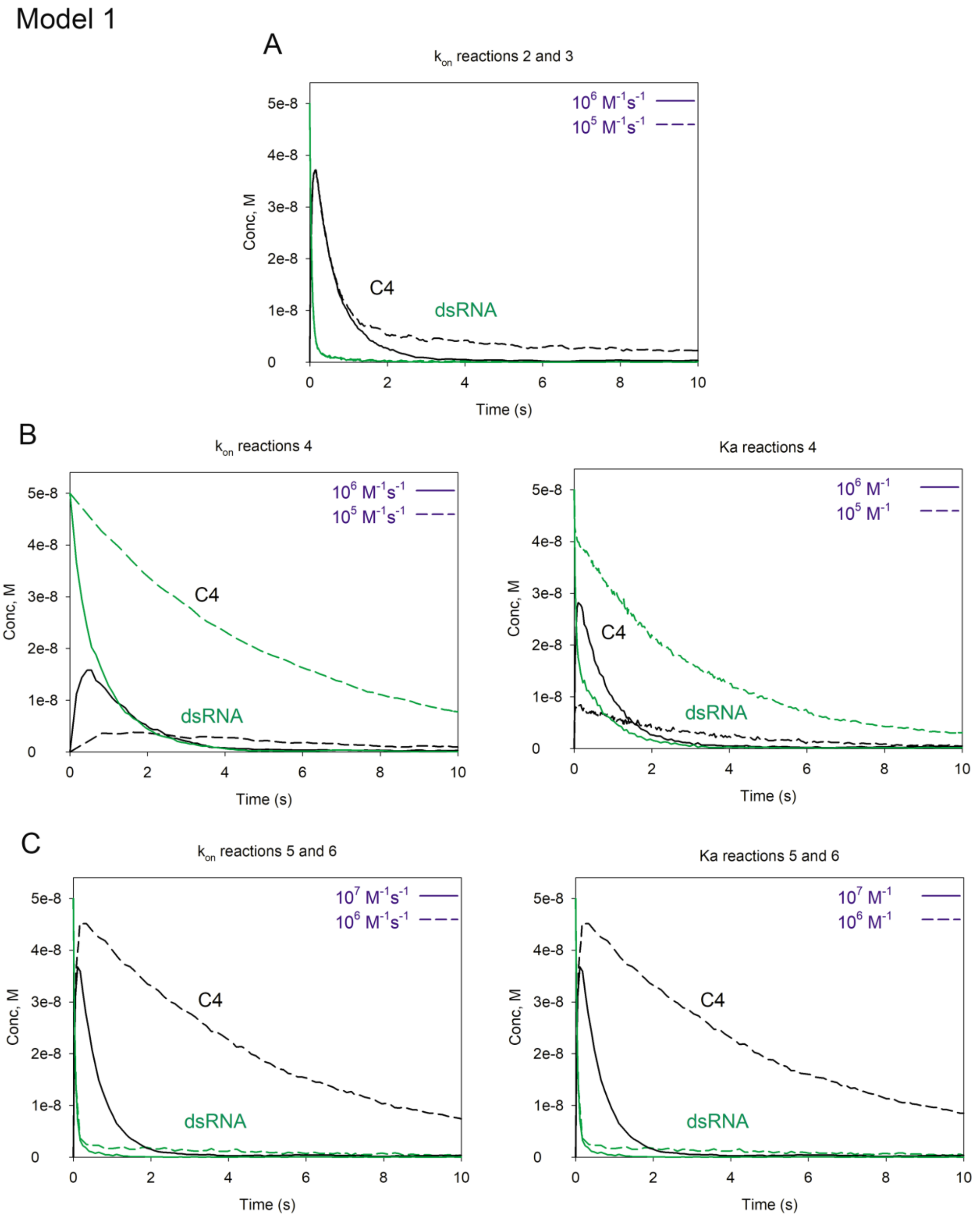


**Figure S14.** Simulations of the reactions progression for the validation of the ranges described in Figure 6A for model 1. A) Effect of the variation of $k_{on}$ in reaction R2 and R3. Note that $k_{on} < 10^6$ $M^{-1}s^{-1}$ makes the reaction too slow to reach equilibrium, violating boundary B4. B) Effect of the variation of $k_{on}$ (left) and $K_a$ (right) in reaction R4. Note that

$k_{on} < 10^6$ $M^{-1}s^{-1}$ or $K_a < 10^6$ $M^{-1}$ make the reaction too slow to reach equilibrium, violating boundary B4. C) Effect of the variation of $k_{on}$ (left) and $K_a$ (right) in reactions R5 and R6. Note that $k_{on} < 10^6$ $M^{-1}s^{-1}$ or $K_a < 10^6$ $M^{-1}$ make the reaction too slow to reach equilibrium, violating boundary B4. For model 1 simulations, we used the following reaction rates: (R1) $k_{on} = 4\times10^{-1}$ $M^{-1}s^{-1}$ and $k_{off} = 8\times10^{-4}$ $s^{-1}$; (R2, R3) $k_{on} = 4\times10^{7}$ $M^{-1}s^{-1}$ and $k_{off} = 1$ $s^{-1}$; (R4) $k_{on} = 1\times10^{7}$ $M^{-1}s^{-1}$ and $k_{off} = 1$ $s^{-1}$; (R5, R6) $k_{on} = 4\times10^{7}$ $M^{-1}s^{-1}$ and $k_{off} = 1$ $s^{-1}$ (red).

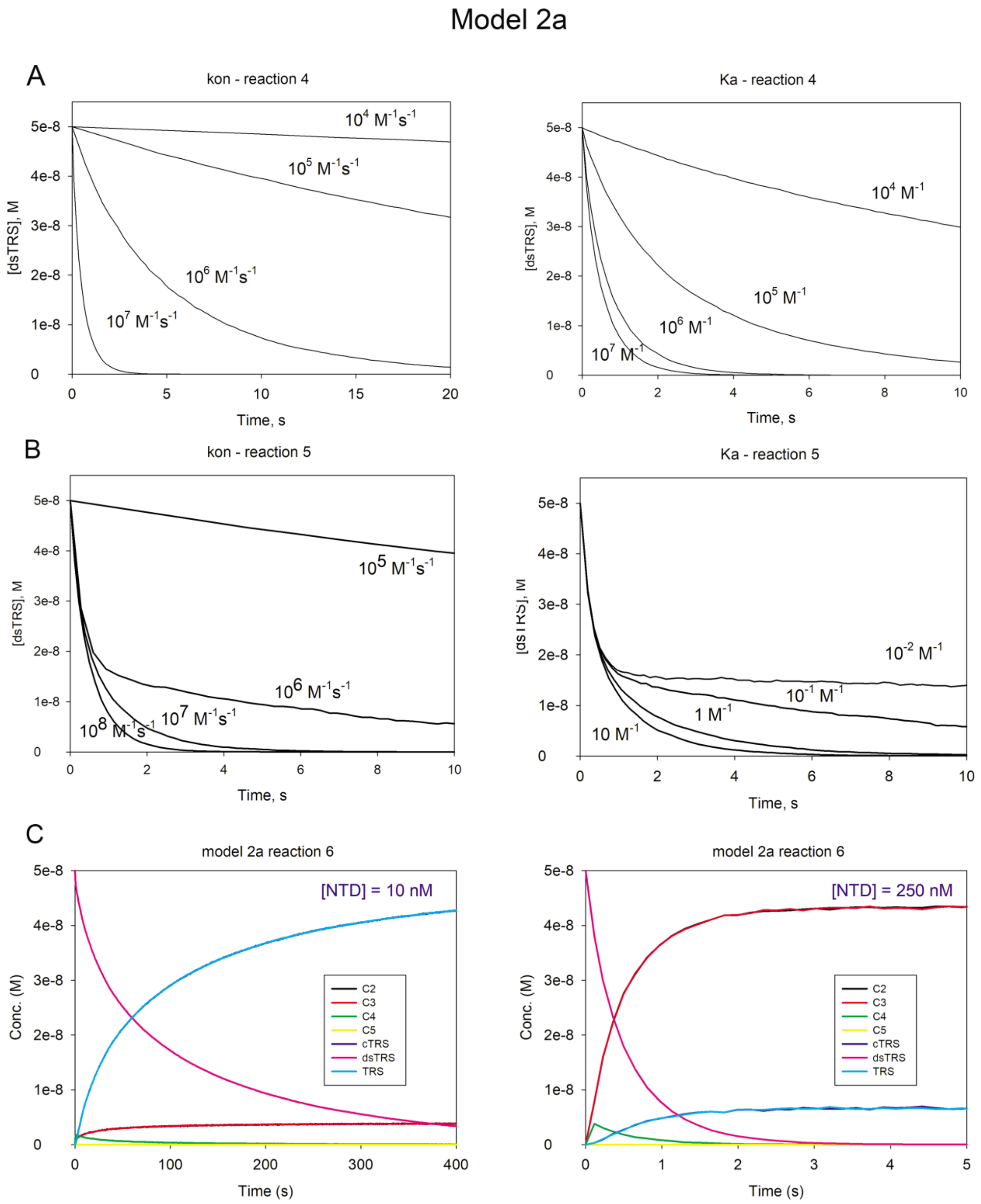


**Figure S15.** Simulations of the reactions progression for the validation of the ranges described in Figure 6A for model 2a. A) Effect of the variation of $k_{on}$ (left) and $K_a$ (right) in reaction R4. Note that $k_{on} < 10^6$ $M^{-1}s^{-1}$ or $K_a < 10^5$ $M^{-1}$ make the reaction too slow to reach equilibrium, violating boundary B4. B) Effect of the variation of $k_{on}$ (left) and $K_a$ (right) in reaction R5. Note that $k_{on} < 10^7$ $M^{-1}s^{-1}$ or $K_a < 1$ $M^{-1}$ make the reaction too slow to reach equilibrium, violating boundary B4. C) Time course of the reaction R6 for each of the components. For model 2a, for $10^7 > K_a > 10^{-6}$ $M^{-1}$, there is never accumulation of C5, resulting in a kinetic of dsRNA melting independent of $k_{on}$ and $k_{off}$ at fixed concentrations of N-NTD. The kinetics changes considerably with the [N-NTD] as showed in the figure.

For values of $K_a > 10^7$ $M^{-1}$ we observed the transition to model 2b with accumulation of C5. Note that for reactions R2 and R3, $K_a$ was determined experimentally ($4\times10^7$ $M^{-1}$). Particularly for model 2a, the kinetic of dsRNA melting is independent of $k_{on}$ and $k_{off}$ of reactions R2 and R3, at fixed concentrations of N-NTD. For model 2a simulations, we used the following reaction rates: (R1) $k_{on} = 4\times10^{-1}$ $M^{-1}s^{-1}$ and $k_{off} = 8\times10^{-4}$ $s^{-1}$; (R2, R3) $k_{on} = 4\times10^7$ $M^{-1}s^{-1}$ and $k_{off} = 1$ $s^{-1}$; (R4) $k_{on} = 1\times10^7$ $M^{-1}s^{-1}$ and $k_{off} = 1$ $s^{-1}$; (R5) $k_{on} = 1\times10^8$ $M^{-1}s^{-1}$ and $k_{off} = 1$ $s^{-1}$ and (R6) $k_{on} = 1\times10^8$ $M^{-1}s^{-1}$ and $k_{off} = 1\times10^{-1}$ $s^{-1}$.

### Model 2b

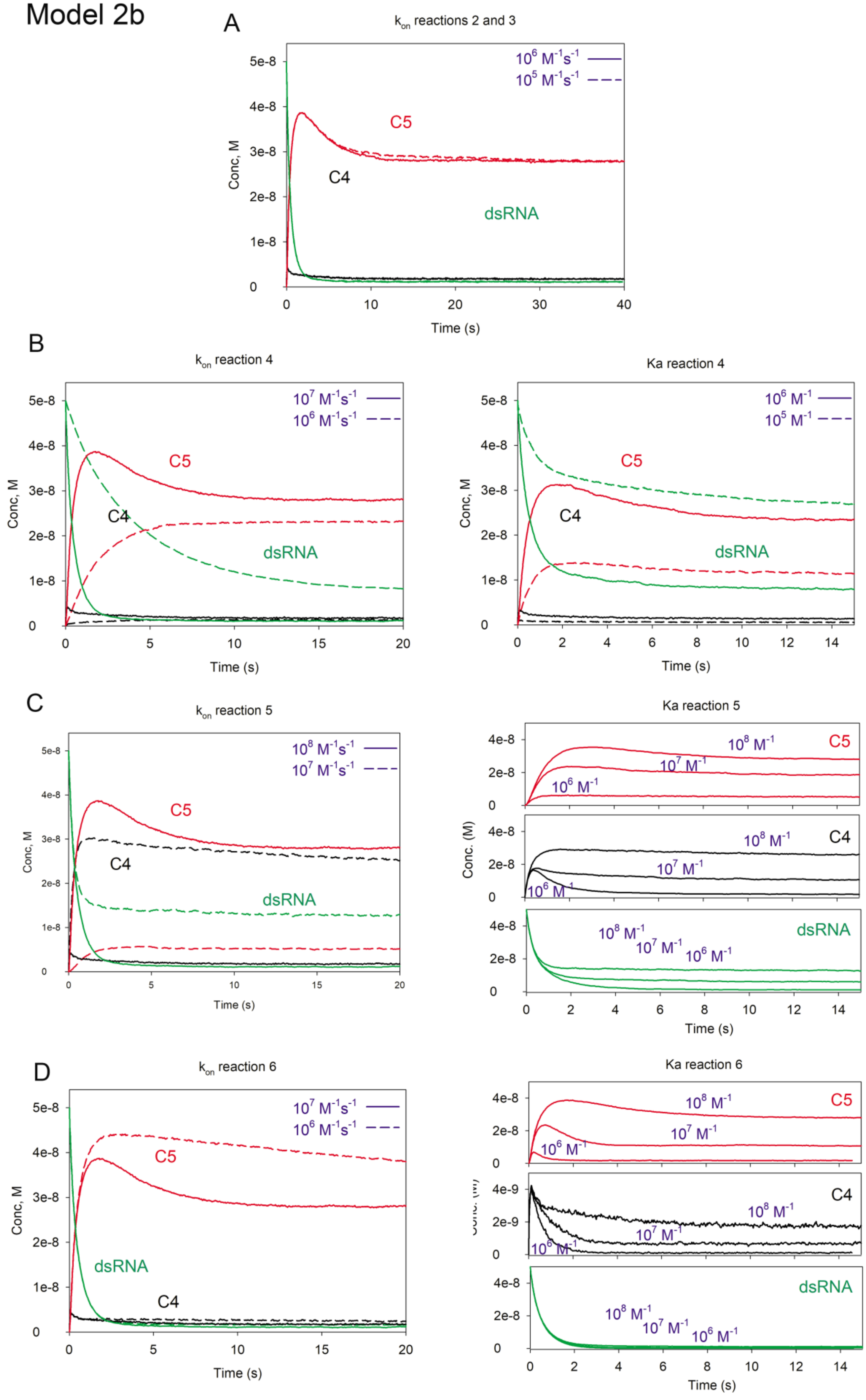
A
$k_{on}$ reactions 2 and 3
$10^6$ $M^{-1}s^{-1}$
$10^5$ $M^{-1}s^{-1}$
C5
C4
dsRNA
Conc, M
Time (s)
B
$k_{on}$ reaction 4
$10^7$ $M^{-1}s^{-1}$
$10^6$ $M^{-1}s^{-1}$
C5
C4
dsRNA
Conc, M
Time (s)
Ka reaction 4
$10^6$ $M^{-1}$
$10^5$ $M^{-1}$
C5
C4
dsRNA
Conc, M
Time (s)
C
$k_{on}$ reaction 5
$10^8$ $M^{-1}s^{-1}$
$10^7$ $M^{-1}s^{-1}$
C5
C4
dsRNA
Conc, M
Time (s)
Ka reaction 5
$10^8$ $M^{-1}$
$10^7$ $M^{-1}$
$10^6$ $M^{-1}$
C5
C4
dsRNA
Conc. (M)
Time (s)
D
$k_{on}$ reaction 6
$10^7$ $M^{-1}s^{-1}$
$10^6$ $M^{-1}s^{-1}$
C5
dsRNA
C4
Conc, M
Time (s)
Ka reaction 6
$10^8$ $M^{-1}$
$10^7$ $M^{-1}$
$10^6$ $M^{-1}$
C5
C4
dsRNA
Time (s)

**Figure S16.** Simulations of the reactions progression for the validation of the ranges described in Figure 6A for model 2b. A) Effect of the variation of $k_{on}$ in reactions R2 and R3. Note that $k_{on} < 10^6$ $M^{-1}s^{-1}$ makes the reaction too slow to reach equilibrium, violating boundary B4. B) Effect of the variation of $k_{on}$ (left) and $K_a$ (right) in reaction R4. Note that $k_{on} < 10^7$ $M^{-1}s^{-1}$ or $K_a < 10^6$ $M^{-1}$ make the reaction too slow to reach equilibrium, violating boundary B4. C) Effect of the variation of $k_{on}$ (left) and $K_a$ (right) in reaction R5. Note that $k_{on} < 10^7$ $M^{-1}s^{-1}$ or $K_a < 10^7$ $M^{-1}$ make the reaction too slow to reach equilibrium, violating boundary B4. D) Effect of the variation of $k_{on}$ (left) and $K_a$ (right) in reaction R6. Note that $k_{on} < 10^6$ $M^{-1}s^{-1}$ or $K_a < 10^7$ $M^{-1}$ make the reaction too slow to reach equilibrium, violating boundary B4. For model 2b simulations, we used the following reaction rates: (R1) $k_{on} = 4\times10^{-1}$ $M^{-1}s^{-1}$ and $k_{off} = 8\times10^{-4}$ $s^{-1}$; (R2, R3) $k_{on} = 4\times10^7$ $M^{-1}s^{-1}$ and $k_{off} = 1$ $s^{-1}$; (R4) $k_{on} = 1\times10^7$ $M^{-1}s^{-1}$ and $k_{off} = 1$ $s^{-1}$; (R5) $k_{on} = 1\times10^8$ $M^{-1}s^{-1}$ and $k_{off} = 1$ $s^{-1}$ and (R6) $k_{on} = 1\times10^8$ $M^{-1}s^{-1}$ and $k_{off} = 1\times10^{-1}$ $s^{-1}$.

#### REFERENCES


1. Baker NA, Sept D, Joseph S, Holst MJ, McCammon JA. 2001. Electrostatics of nanosystems: Application to microtubules and the ribosome. Proc Natl Acad Sci U S A 98:10037–10041.
2. Dolinsky TJ, Czodrowski P, Li H, Nielsen JE, Jensen JH, Klebe G, Baker NA. 2007. PDB2PQR: expanding and upgrading automated preparation of biomolecular structures for molecular simulations. Nucleic Acids Res 35:W522-5.
3. Olsson MHM, SØndergaard CR, Rostkowski M, Jensen JH. 2011. PROPKA3: Consistent treatment of internal and surface residues in empirical p K a predictions. J Chem Theory Comput 7:525–537.
4. Delano WL. 2002. The PyMOL Molecular Graphics System. DeLano Scientific, San Carlos, CA, USA. San Carlos, CA, USA.
5. Lu X-J, Olson WK. 2003. 3DNA: a software package for the analysis, rebuilding and visualization of three-dimensional nucleic acid structures. Nucleic Acids Res 31:5108–5121.
